# Quiet wakefulness: The influence of intraperitoneal and intranasal oxytocin on sleep-wake behaviour and neurophysiology in rats

**DOI:** 10.1101/2022.11.23.514802

**Authors:** Joel S Raymond, Nicholas A Everett, Anand Gururajan, Michael T Bowen

## Abstract

**Introduction:** Exogenous administration of the neuropeptide oxytocin exerts diverse effects on various neurobehavioural processes, including sleep and wakefulness. Since oxytocin can enhance attention to social and fear-related environmental cues, it should promote arousal and wakefulness. However, as oxytocin can attenuate stress, reduce activity, and elicit anxiolysis, oxytocin might also prime the brain for rest and promote sleep. At present, little research has comprehensively characterised the neuropsychopharmacology of oxytocin-induced effects on sleep-wake behaviour and no reconciliation of these two competing hypotheses has been proposed.

**Methods:** This study explored the effects of oxytocin on sleep-wake outcomes using radiotelemetry-based polysomnography in adult male and female Wistar rats. Oxytocin was administered via intraperitoneal (i.p.; 0.1, 0.3 and 1 mg·kg^-1^) and intranasal (i.n.; 0.06, 1, 3 mg·kg^-1^) routes. Caffeine (i.p. and i.n.; 10 mg·kg^-1^) was administered as a wake-promoting positive control. To ascertain mechanism of action, pre-treatment experiments with the oxytocin receptor (OXTR) antagonist L-368,899 (i.p.; 5 mg·kg^-1^) followed by oxytocin (i.p.; 1 mg·kg^-1^) were also conducted.

**Results:** In both male and female rats, i.p. oxytocin promoted quiet wakefulness at the cost of suppressing active wakefulness, NREM and REM sleep. Several i.p. oxytocin-induced sleep-wake effects were mediated by OXTR binding. In contrast, i.n. oxytocin did not alter most sleep-wake outcomes at any dose tested. Both i.p. and i.n. caffeine demonstrated wake-promoting effects.

**Conclusions:** These findings help reconcile competing hypotheses of oxytocin-induced effects on sleep-wake behaviour: i.p. oxytocin promotes quiet wakefulness—a state of restful environmental awareness compatible with both oxytocin’s anxiolytic effects and its enhancement of processing complex stimuli.

## Introduction

The neuropeptide oxytocin exerts diverse effects on numerous domains of mammalian neurobiology and behaviour ^1^, including sleep and wakefulness ^2^. Endogenous oxytocin is synthesised in the paraventricular (PVN) and supraoptic nuclei (SON) of the hypothalamus, and is released into circulation via the neurohypophysis, and centrally via axonal projections to various forebrain, midbrain and hindbrain regions ^3, 4^. As both oxytocinergic projections and oxytocin receptor (OXTR) expression have been found in various hypothalamic and pontine sleep-wake regulatory brain regions ^1, 4^, the fundamental neural circuitry exists for oxytocin to impact sleep and wakefulness. However, current understanding of oxytocin’s influence on sleep-wake behaviour and neurophysiology is limited ^2^.

Theoretically, it is difficult to hypothesise what impact oxytocin will exert on sleep-wake outcomes as two seemingly contradictory hypotheses emerge. From an *environmental awareness-arousal* perspective, oxytocin can orient attention towards environmental cues critical for survival and adaptation ^5^, both social ^6, 7^ and fear-related^8, 9^. Enhanced attention likely relies on sufficiently enhanced physiological arousal ^10^, so hypothetically oxytocin should promote arousal and consequently, promote wakefulness. In contrast, from a *stress attenuation-quiescence* perspective, oxytocin can reduce stress and anxiety ^11–13^, locomotor activity ^14, 15^, pain sensitivity ^16, 17^, heart rate ^18^, and aggression ^19^. Taken together, this suggests that oxytocin physiologically and psychologically primes the brain and body for rest, which in turn should— hypothetically—promote sleep.

Empirically, based on our recent systematic review ^2^, exogenous oxytocin appears to exert mixed sleep-wake effects ranging from promoting wakefulness ^20, 21^ to promoting sleep ^22^ to exerting no sleep-wake effects ^23, 24^ across preclinical studies. However, when factors of dose and route of administration were considered, a relatively clear wake-promoting effect of oxytocin emerged. Additionally, a lack of studies investigating potential sex differences in oxytocin-induced sleep-wake effects was highlighted. Hence, the current study aimed to address the following research questions: (1) what effects does peripherally administered oxytocin exert on sleep-wake behaviour and neurophysiology; (2) do any effects depend on dose, route of administration, or biological sex; and (3) are any observed effects mediated by the OXTR? Based on preclinical evidence from our systematic review ^2^, we hypothesised that oxytocin would exert dose-dependent wake-promoting effects.

## Method

An abridged method section is presented below; for complete methodological details, see ‘Method’ section in supplemental material.

### Animals and housing

Eight-week-old male and female Wistar rats (ARC, WA, Australia) were pair-housed in filter-top cages (58 x 38 x 20 cm; Able Scientific) containing corn cob bedding material (Bed-o’ Cobs, The Andersons), environmental enrichment, and *ad libitum* access to standard rodent chow and water. All experiments were conducted within a specific pathogen free (SPF) facility and rats were housed in a temperature- and humidity-controlled room (22 ± 0.5°C; 50-60%) under a reverse light cycle (12L:12D; lights on at 1400, defined as ZT0). At transitions between light phases, illumination was slowly transitioned between 0 and 500 lux over a 16-min period. All experiments were conducted in line with the *Australian code for the care and use of animals for scientific purposes (8^th^ edition, 2013)* and were approved by the Animal Ethics Committee at The University of Sydney (AEC number: 2019/1615).

### Drug preparation

Oxytocin (China Peptide, China) and caffeine (anhydrous; AK Scientific Inc., USA) were dissolved in physiological saline (0.9% w/v). The non-peptidergic OXTR antagonist L-368,899 hydrochloride (Santa Cruz Biotechnology, Texas) was dissolved in dimethylsulfoxide (5% v/v), Tween 80 (5% v/v), and saline (90% v/v).

### Intraperitoneal drug administration

All drugs administered via intraperitoneal (i.p.) injection were administered at an injection volume of 1 mL·kg^-1^ body weight. Oxytocin was administered at light onset (ZT0) at a range of doses (0.1, 0.3, and 1 mg·kg^-1^ b.w.). These doses were selected to cover doses of oxytocin that do (1 mg·kg^-1^) and do not (0.3 and 0.1 mg·kg^-1^) supress gross locomotor activity in rats ^14^. Caffeine was administered at ZT0 at a dose of 10 mg·kg^-1^ as a positive control, based on previous research demonstrating potent wake-promoting effects at this dose in rats ^25, 26^. OXTR antagonist (L-368,899, 5 mg·kg^-1^) pre-treatment was administered 15-min prior to light onset (ZT23.75). L-368,899 was chosen as it is centrally penetrant and preferentially binds to OXTRs ^27, 28^. The dose was chosen based on its ability to block the stress-attenuating effects of 1 mg·kg^-1^ oxytocin in rats ^29^.

### Intranasal drug administration

Oxytocin (0.06, 1, and 3 mg·kg^-1^) and caffeine (10 mg·kg^-1^) were administered intranasally (i.n.) according to methods adapted from Lukas and Neumann ^30^. The 0.06 mg·kg^-1^ dose represents an approximately four-to-five-fold higher dose than the highest i.n. dose administered in human clinical research 80 IU; ^31^; based on allometric scaling using the FDA guidance for conversion between animal and human doses Food and Drug ^32^, the human equivalent dose of 0.06 mg·kg^-1^ in a rat is ∼348 IU for a 60 kg human. This dose was selected to investigate whether doses closer to those administered in clinical research would impact sleep-wake behaviour. The 1 and 3 mg·kg^-1^ doses were chosen to facilitate comparisons between i.p. and i.n. routes of administration; the higher dose was based on a previous study demonstrating that i.p. oxytocin elicited approximately two-fold greater increases in plasma and central oxytocin levels compared to the same dose administered i.n. ^33^.

### Radiotelemetry probe implantation surgery

Briefly, after induction and administration of pre-operative analgesics, rats were surgically implanted with wireless radiotelemetry probes capable of polysomnographic (PSG)—electrocorticographic (ECoG) and electromyographic (EMG)—recording (HD-X02, Data Sciences International Inc.). The telemetry probe was implanted subcutaneously under the right flank, ECoG wires were secured around two screws contacting the dura at fronto-parietal locations (measured from bregma: (1) anterior/posterior: +2 mm, lateral: +1.5 mm; (2) anterior/posterior: –7 mm, lateral: –1.5 mm), and EMG wires were threaded through and secured to trapezius muscle. Rats received post-operative analgesia and were allowed at least 14 days recovery prior to experimentation.

### Polysomnographic recordings

Rats were placed individually into telemetry cages (43 x 26 x 12 cm, Techniplast, Italy) located above telemetry receiver plates (RPC-1, Data Sciences International Inc.). Habituation to the telemetry recording cages and procedures was conducted over three 6-hour baseline PSG recordings performed prior to testing to ensure proper probe functioning and reception by telemetry receiver plates. For all experiments except Experiment 2, rats were placed into recording cages at least 1 hour prior to light onset and recordings were started by 1300 (ZT23). Due to pre-treatment with antagonist L-368,899 for Experiment 2, rats were placed into recording cages at least 1.25 hour prior to light onset and recordings were started by 1245 (ZT22.75). At light onset (ZT0), rats were removed from the telemetry cage, administered an i.p. injection or i.n. application, placed back in the cage, left to behave freely for 6 hours (until ZT6), and then returned to their home cages (Figure 1A). For female rats, oestrus phase determination was conducted according to methods adapted from Marcondes, Bianchi and Tanno ^34^, immediately following the termination of PSG recording sessions (ZT6-6.5).

**Figure 1.**
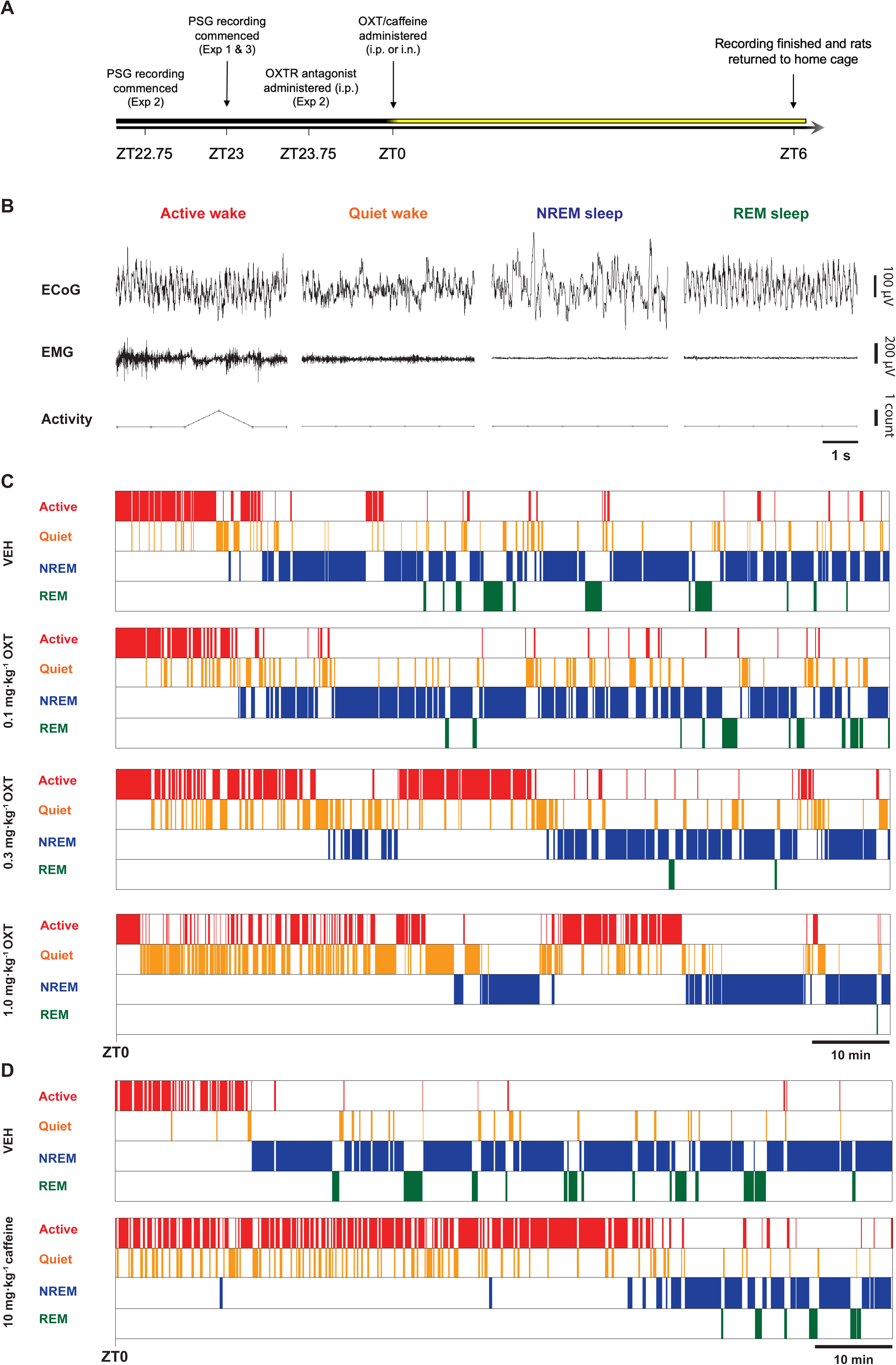
Overview of polysomnographic session timeline, sleep-wake scoring procedure and representative hypnograms from Experiment 1. (A) Experimental schedule outlining the timing of procedures undertaken during recording sessions. The gradient and colour of the bar represent the circadian phase and intensity of light conditions during the shift from the dark phase to the light phase. i.n.: intranasal; i.p.: intraperitoneal; OXT: oxytocin; OXTR: oxytocin receptor; PSG: polysomnography; ZT: zeitgeber, timing relative to light onset (ZT0). (B) Representative polysomnography traces (5-s duration) of each sleep-wake state category used for scoring. ECoG: electrocorticography; EMG: electromyography. (C) Representative hypnograms depicting sleep-wake state across 0-120 min of recording (starting at ZT0) for each dose of i.p. oxytocin administered (0, 0.1, 0.3 and 1 mg·kg^-1^). (D) Representative hypnograms depicting sleep-wake state across 0-120 min of recording (starting at ZT0) for i.p. caffeine (0, 10 mg·kg^-1^). Hypnograms for (C) and (D) were constructed using data from the subject with the median effect size of i.p. oxytocin 1 mg·kg^-1^ on %QW 0-90 min.

### Experimental design

All experiments were conducted across consecutive recording sessions using repeated-measures, counterbalanced designs to minimise inter-subject variability and increase statistical power. In all designs, dose sequences were generated using William’s Latin Square designs that control for first-order carryover effects ^35^, and rats were randomised to dose sequences using a random number generator, with the caveat that pair-housed rats could not be assigned to the same dose sequence. There was at least 90-h washout before the next recording session following oxytocin or L-368,899 administration, and 7 days following caffeine administration, which demonstrates more marked and prolonged effects on sleep ^20, 26^. Hence, the caffeine condition was not incorporated into oxytocin dose-response William’s Latin square designs in Experiment 1 to preserve a consistent washout period between recording sessions.

### Oxytocin i.p. dose-response (Experiments 1A and 1B)

This experiment characterised the effects of i.p. oxytocin (VEH, 0.1, 0.3, 1 mg·kg^-1^; Experiment 1A) and caffeine (VEH and 10 mg·kg^-1^; Experiment 1B) on sleep-wake outcomes. Male and female rats (*N* = 16; *n* = 8 per sex) were run as separate cohorts to avoid potential interference of opposite-sex pheromones in sleep-wake behaviour and physiology during PSG recording sessions reviewed by ^36^.

### Oxytocin i.p. and OXTR antagonism (Experiment 2)

This experiment explored whether i.p. oxytocin-induced effects on sleep-wake outcomes are mediated by the OXTR. Since no significant sex by dose interactions were apparent during Experiment 1 (see Supplemental Material Table S1), only female rats (*n* = 4) were used for Experiment 2. The following conditions examined whether an OXTR antagonist inhibited oxytocin effects on sleep-wake outcomes: (i) VEH + VEH; (ii) VEH + oxytocin (1 mg·kg^-1^); (iii) L-368,899 (5 mg·kg^-1^) + VEH; (iv) L-368,899 (5 mg·kg^-1^) + oxytocin (1 mg·kg^-1^).

### Oxytocin i.n. dose-response (Experiments 3A and 3B)

Experiment 3A characterised the dose-dependent effects of i.n. oxytocin (0, 0.06, 1 mg·kg^-1^ in 20 μL), and Experiment 3B a higher dose of i.n. oxytocin (3 mg·kg^-1^ in 20 μL) and caffeine (10 mg·kg^-1^ in 20 μL), on sleep-wake outcomes. The goal was to examine whether i.n. oxytocin would recapitulate the effects of i.p. oxytocin observed in Experiment 1. As with Experiment 2, only female rats were used: *n* = 6 for Experiment 3A and *n* = 5 for Experiment 3B.

### Data acquisition, processing, and analysis

Polysomnographic recording data (ECoG, EMG, body temperature and activity) were acquired using Ponemah software (Version 6.41, Data Sciences International). NeuroScore (Version 3.2.1, Data Sciences International) was used to score sleep-wake states for each 10-s epoch of time: active wake (AW), quiet wake (QW), non-REM sleep (NREMS) and REM sleep (REMS). Pre-REM sleep, the transition state between NREMS and REMS characterised by high amplitude EEG activity within the theta and alpha power bands ^37^, was scored as NREMS.

ECoG and EMG signals were filtered prior to scoring: a band-pass filter (0.1-80 Hz) and high-pass filter (> 0.1 Hz) were applied to the ECoG and EMG signal Data Sciences ^38^, respectively. Sleep scoring was conducted manually by a blinded experimenter via visual examination of various raw and derived signals: power spectral density ‘periodogram’ of ECoG signal (Fast Fourier Transform; 0-25 Hz), theta:delta ratio, ECoG trace, EMG trace, activity counts, and sleep-wake state of previous epoch.

Briefly, scoring criteria adapted from ^39^ for each sleep-wake state were as follows (see Figure 1B for representative traces):

- *Active wake*: (ECoG) low amplitude, low synchrony, high frequency; (EMG) high amplitude, irregular tone; (Activity) locomotion and movement present
- *Quiet wake*: (ECoG) low amplitude, low synchrony, moderate-high frequency relative to AW; (EMG) moderate amplitude relative to AW, regular tone; (Activity) no locomotion and minimal movement
- *NREM sleep*: (ECoG) moderate-high amplitude, high synchrony, very low-low frequency; (Periodogram) high power density within the delta frequency band (0-4 Hz); (EMG) low amplitude, regular tone; (Activity) no locomotion or movement
- *REM sleep*: (ECoG) low amplitude, high frequency, ‘saw-tooth’ profile; (Periodogram) high power density within the theta frequency band (4-8 Hz) and high theta:delta ratio; (EMG) low amplitude, regular tone; (Activity) no locomotion or movement
- *Artefact*: epochs were scored as artefacts and excluded from analysis if no signals (ECoG, EMG, activity etc.) were available during the epoch to indicate sleep-wake state.

After sleep scoring was completed, the following sleep architectural outcomes of interest were extracted: sleep onset latency (min), REM sleep onset latency (min), proportion of total time spent in each sleep state (%), bout frequency of each sleep state, mean duration of bout of each sleep state (min), and mean body temperature (°C). These 7 hours of PSG data (1-h prior to oxytocin administration during last hour of dark phase and 6 hours post-administration during light phase) were then compiled into 30-min bins for statistical analysis. Sample sizes for each experiment and experimental condition are detailed in figure legends; some subjects were excluded due to surgical complications following probe implantation, while some data is absent due to issues with dropout during telemetry recordings. For complete details on subject attrition and data exclusion, see Supplemental materials.

ECoG power spectral density (PSD) outcomes during QW, NREMS, and REMS were extracted for each 1-Hz frequency band from 0-25 Hz using Discrete Fourier Transform (DFT). Artefact detection, both automated and manual, was conducted and those epochs containing artefacts were excluded Data Sciences ^38^. As large inter-individual differences existed in ECoG spectral outcomes, all subjects’ data were normalised by expressing values as a proportion of an outcome during the corresponding VEH condition of each respective experiment ^20^. For further details on PSD data analysis, see Supplemental materials.

As sleep outcomes appear to be predominantly influenced by the oestrus cycle during proestrus and oestrus phases ^40, 41^, oestrus stage was delineated into two categories: either proestrus/oestrus or metoestrus/dioestrus.

### Statistical analysis

Linear mixed effects models (LMMs) were constructed for all sleep architectural outcomes with *Dose*, *Time,* and *Dose* x *Time* as within-subjects fixed effects factors. LMMs were chosen to avoid having to exclude subjects with missing data. A compound symmetry covariance matrix structure was employed and fitted using restricted maximum likelihood estimation. Greenhouse-Geisser corrections were applied to adjust for violations of sphericity. If random effects were zero, they were removed from the model and a simpler model was fitted.

For sleep architectural outcomes, the first hour of data (dark phase; pre-administration) were analysed separately from the following 6 hours (light phase; post-administration) to test for baseline differences in outcomes. As part of the LMM analysis, trend analysis was conducted on *Dose* to determine if a linear dose-dependent effect was present. Based on the pharmacokinetics of acute peripherally administered oxytocin (t_1/2_ = < 5 min ^42^) and *post hoc* visual inspection of the sleep architectural outcomes, average values were calculated for periods of the peak effect for each outcome—AW: 0-30 min and 30-180 min, QW: 0-90 min, NREMS: 0-90 min, REMS: 30-180 min, and body temperature: 0-120 min. LMM and trend analyses were conducted on these composite outcomes with *Dose* as a fixed effect.

For ECoG PSD outcomes, normalised average values (%) for each 1-Hz frequency band were transformed using a logarithmic (log10) transformation, and then compared to the baseline (VEH) value using a two-tailed one-sample t-test. For oestrus phase data, differences in the proportion of rats in proestrus/oestrus and metoestrus/dioestrus phases during each experiment between each dose and VEH were analysed using two-sided Fisher’s Exact tests. Fisher’s least significant difference (LSD) test was used to control Type I error rate. Statistical analyses were conducted using GraphPad Prism (Version 9.3.1). The level of significance for all tests was *p* < .05.

## Results

For Experiment 1, data from male and female rats were combined for analyses based on the lack of significant *Sex* x *Dose* interaction effects observed across all proportion of time spent in sleep-wake state outcomes for periods of peak effect for Experiment 1a (for more details, see supplemental material Table S1).

### Experiment 1 – Effects of i.p. oxytocin dose range and i.p. caffeine on sleep-wake outcomes

Figure 1 depicts the timeline of sleep recording sessions (1A), representative PSG traces for each sleep-wake state (1B), and representative hypnograms of results from Experiment 1 illustrating the temporal nature of oxytocin- and caffeine-induced effects on sleep-wake state (1C and D). Summarised statistical results for Experiments 1a and 1b are presented in Tables 1 and S2.

There were no significant differences in the proportion of female rats in different oestrus phases between dose conditions (see Tables S3 and S4). Additionally, there were no significant main effects of dose across %sleep-wake state outcomes at baseline (see Tables S5 and S6).

#### Wake outcomes

The effects of i.p. oxytocin (0, 0.1, 0.3 and 1 mg·kg^-1^) and caffeine (0, 10 mg·kg^-1^) on wake outcomes are displayed in Figure 2 and corresponding statistical outcomes are reported in Tables 1 and S2.

**Figure 2.**
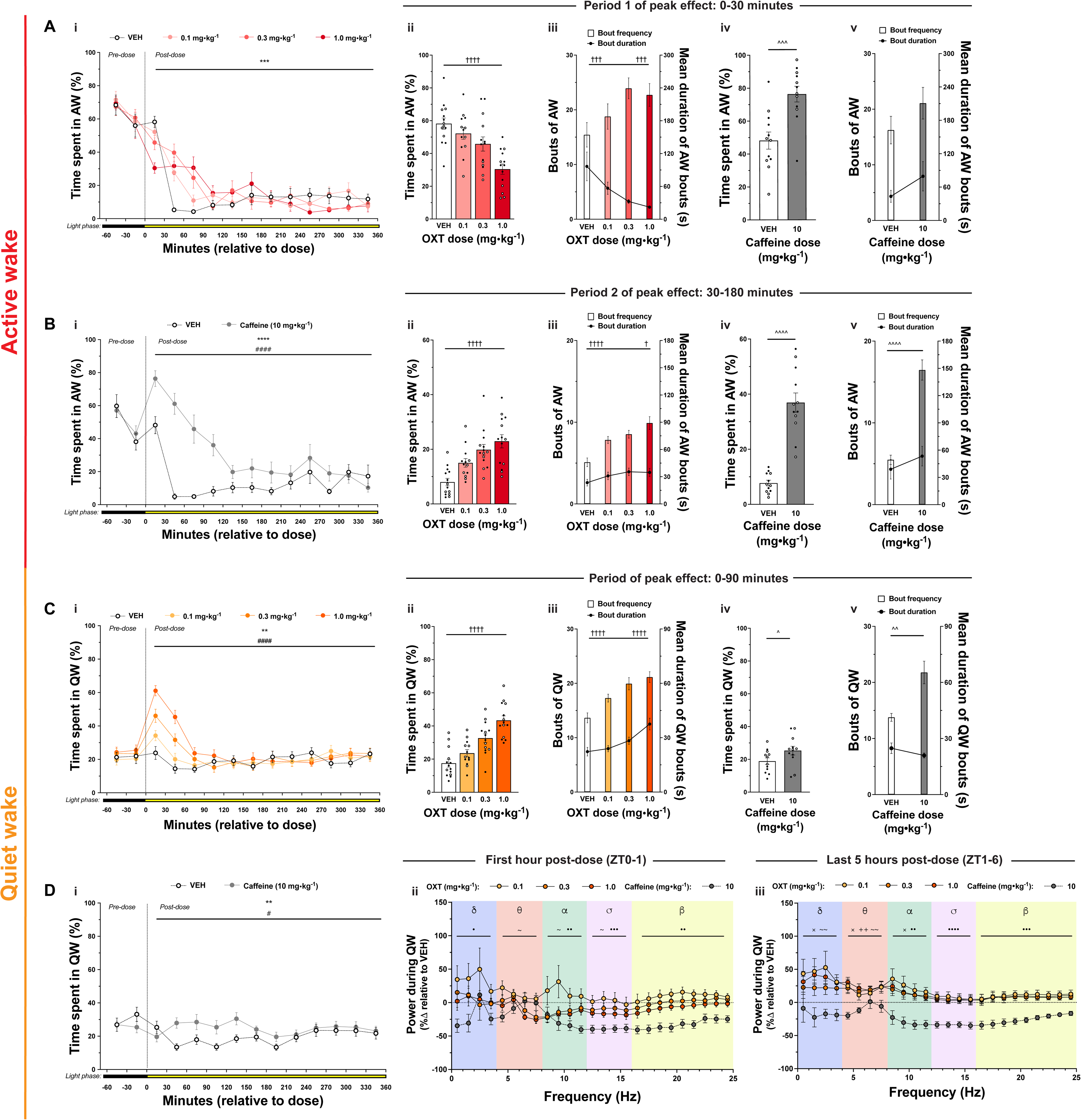
Effects of i.p. oxytocin and i.p. caffeine on wake outcomes. *Oxytocin effects.* Influence of i.p. oxytocin (0, 0.1, 0.3, and 1 mg·kg^-1^) on %AW across entire 7-hour recording session (**Ai**), %AW during first period of peak effect (0-30 min) (**Aii**), AW bout frequency and mean AW bout duration during first period of peak effect (0-30 min) (**Aiii**), %AW during second period of peak effect (30-180 min) (**Bii**), AW bout frequency and mean AW bout duration during second period of peak effect (30-180 min) (**Biii**), %QW across entire 7-hour recording session (**Ci**), %QW during first period of peak effect (0-90 min) (**Cii**), and QW bout frequency and mean QW bout duration during first period of peak effect (0-90 min) (**Ciii**). *Caffeine effects.* Influence of i.p. caffeine (0 and 10 mg·kg^-1^) on %AW across entire 7-hour recording session (**Bi**),%AW during first period of peak effect (0-30 min) (**Aiv**), AW bout frequency and mean AW bout duration during first period of peak effect (0-30 min) (**Av**), %AW during second period of peak effect (30-180 min) (**Biv**), AW bout frequency and mean AW bout duration during second period of peak effect (30-180 min) (**Bv**), %QW across entire 7-hour recording session (**Di**), %QW during period of peak effect (0-90 min) (**Civ**), and QW bout frequency and mean QW bout duration during period of peak effect (0-90 min) (**Cv**). *Power spectral density effects.* Influence of i.p. oxytocin and caffeine on ECoG power spectral density from 0-1 h (**Dii**) and 1-6 h during QW (**Diii**). Sample sizes for oxytocin dose-response are *n* = 6 males (for all doses) and *n* = 8 females for all doses except 0.1 mg·kg^-1^ which is *n* = 7. Sample sizes for caffeine positive control are *n* = 6 males and *n* = 6 females for all doses. Data represent mean values ± S.E.M., individual data points represent individual subject data (open circles – males; closed circles – females). For % time spent in sleep-wake state graphs, data points represent values for 30-min bins; values pertain to the bin of time defined by the x-axis timepoints to the immediate left and right of the data point. Dose was administered at ZT0 and colour of bar below x-axis signifies light cycle phase at each time point: black – dark (active) phase; yellow – light (rest) phase. For ECoG PSD figures, data represent mean percentage change in ECoG PSD (0.1-25 Hz) ± S.E.M. Statistical significance is indicated by the following symbols: * – dose main effect; # – dose x time interaction effect; † – linear trend contrast; ^ pairwise comparison; × – 0.1 mg·kg^-1^ oxytocin vs VEH; + – 0.3 mg·kg^-1^ oxytocin vs VEH; ∼ – 1 mg·kg^-1^ oxytocin vs VEH; • – 10 mg·kg^-1^ caffeine vs VEH. For graphs with two y-axes, the alignment of the significance symbol represents which outcome the significance refers to: left – bout frequency; right – mean bout duration. Level of statistical significance is indicated by the number of symbols: one – *p* < .05, two – *p* < .01, three – *p* < .001, four – *p* < .0001.

#### Active wake

For %AW, over the 360 min post-dose period, there was an oxytocin *Dose* x *Time* interaction (Figure 2Ai). During 0-30 min, oxytocin dose-dependently reduced %AW (Figure 2Aii); this effect was produced through a dose-dependent reduction in mean AW bout duration, which was, interestingly, accompanied by a dose-dependent *increase* in bouts of AW (Figure 2Aiii). In contrast, during 30-180 min, oxytocin dose-dependently increased %AW (Figure 2Bii) due to a dose-dependent increase in AW bouts and mean AW bout duration (Figure 2Biii).

Overall, caffeine (i.p.; 10 mg·kg^-1^) increased %AW and there was a *Dose* x *Time* interaction (Figure 2Bi). During 0-30 min, caffeine increased %AW (Figure 2Aiv), however it was unclear whether this was due to increased bouts of AW or increased mean AW bout duration (Figure 2Av). During 30-180 min, caffeine also increased %AW (Figure 2Biv), through increased AW bouts but not mean AW bout duration (Figure 2Bv).

#### Quiet wake

Overall, oxytocin impacted %QW and this changed over time during the sleep recording (Figure 2Ci). During 0-90 min, oxytocin dose-dependently increased %QW (Figure 2Cii) by increasing both QW bouts and mean QW bout duration (Figure 2Ciii). Overall, caffeine increased %QW and this differed over the course of the sleep recording (Figure 2Di). During 0-90 min, caffeine increased %QW (Figure 2Civ), which was due to increased QW bouts but not mean QW bout duration (Figure 2Cv).

During the 1^st^ hour post-dose, relative to vehicle, 1 mg·kg^-1^ oxytocin reduced ECoG PSD in the theta, alpha and sigma frequency bands, while caffeine reduced ECoG PSD in the delta, alpha, sigma, and beta frequency bands (Figure 2Dii). In contrast, during the last 5 hours of recording, oxytocin increased ECoG PSD in the delta (0.1 and 1 mg·kg^-1^), theta (all doses), and alpha (0.1 mg·kg^-1^) frequency bands, whereas caffeine reduced ECoG PSD within the alpha, sigma, and beta frequency bands relative to vehicle (Figure 2Diii).

#### Sleep outcomes

##### NREM sleep

Overall, for %NREMS, there was an oxytocin *Dose* x *Time* interaction (Figure 3Ai). During 0-90 min, oxytocin dose-dependently reduced %NREMS (Figure 3Aii), due to both reduced bouts of NREMS and reduced mean NREMS bout duration (Figure 3Aiii). However, there was no effect of oxytocin on sleep onset latency (Figure 3Aiv).

**Figure 3.**
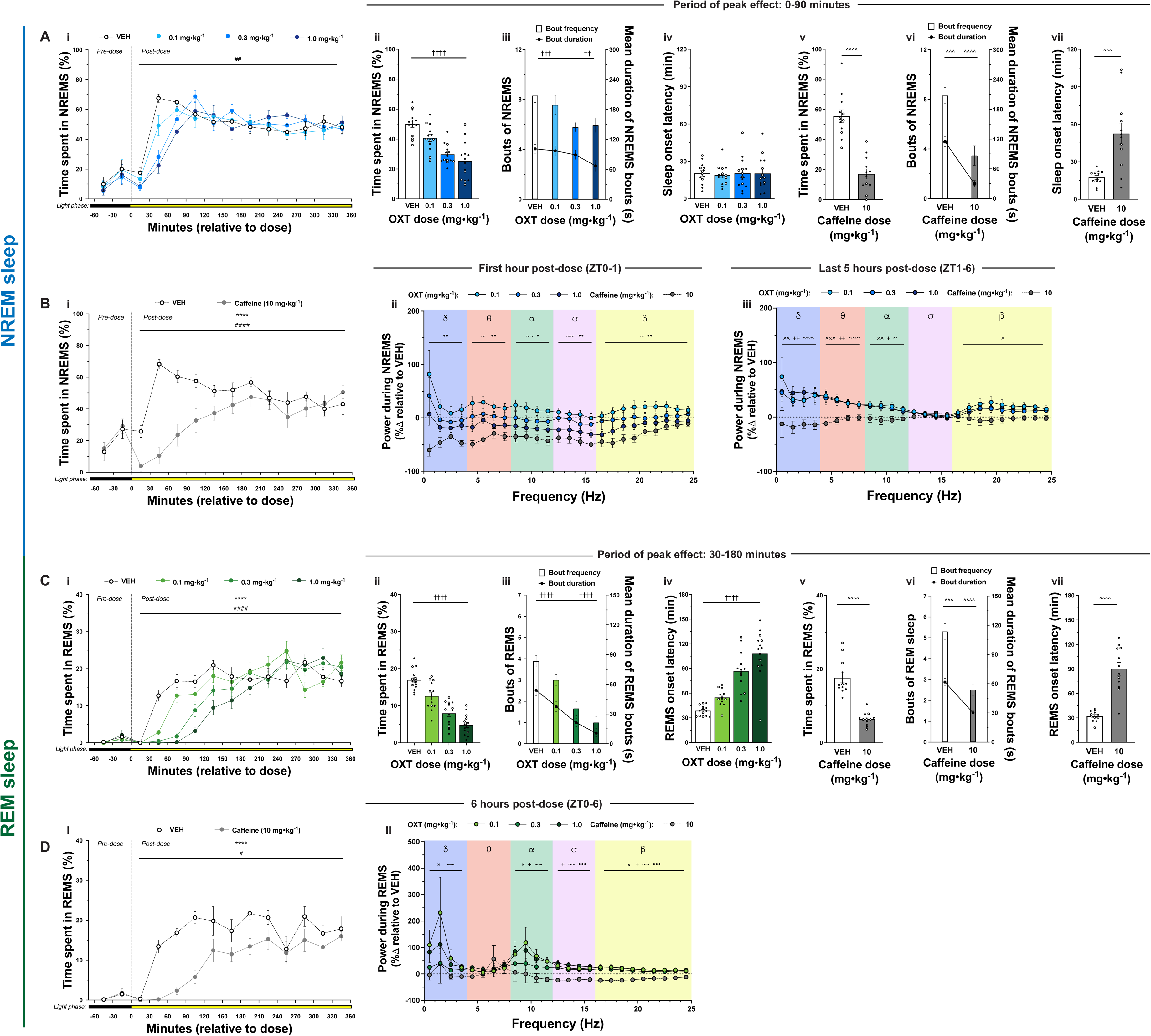
Effects of i.p. oxytocin and i.p. caffeine on sleep outcomes. *Oxytocin effects.* Influence of i.p. oxytocin (0, 0.1, 0.3, and 1 mg·kg^-1^) on %NREMS sleep across entire 7-hour recording session **(Ai)**, %NREMS during period of peak effect (0-90 min) **(Aii)**, NREMS bout frequency and mean NREMS bout duration during period of peak effect (0-90 min) **(Aiii)**, sleep onset latency **(Aiv)**, %REMS across entire 7-hour recording session **(Ci)**, %REM sleep during period of peak effect (30-180 min) **(Cii)**, REMS bout frequency and mean REMS bout duration during period of peak effect (30-180 min) (**Ciii)**, and REMS onset latency (**Civ)**. *Caffeine effects.* Influence of i.p. caffeine (0 and 10 mg·kg^-1^) on %NREMS across entire 7-hour recording session (**Bi)**, %NREMS during period of peak effect (0-90 min) (**Av)**, NREMS bout frequency and mean bout duration during period of peak effect (0-90 min) **(Avi)**, sleep onset latency **(Avii)**, %REM sleep across entire 7-hour recording session **(Di**), %REM sleep during period of peak effect (30-180 min) **(Cv)**, REMS bout frequency and mean bout duration during period of peak effect (30-180 min) **(Cvi)**, and REMS onset latency **(Cvii)**. *Power spectral density effects.* Influence of i.p. oxytocin dose range and caffeine on ECoG power spectral density from 0-1 h and 1-6 h during NREMS (**Bi** and **Bii**, respectively) and 0-6 h during REMS **(Dii)**. Sample sizes for oxytocin dose-response are *n* = 6 males (for all doses) and *n* = 8 females for all doses except 0.1 mg·kg^-1^ which is *n* = 7. Sample sizes for caffeine positive control are *n* = 6 for males and females for all doses. Data represent mean values ± S.E.M., individual data points represent individual subject data (open circles – males; closed circles – females). For % time spent in sleep-wake state graphs, data points represent values for 30-min bins; values pertain to the bin of time defined by the x-axis timepoints to the immediate left and right of the data point. Dose was administered at ZT0 and colour of bar below x-axis signifies light cycle phase at each time point: black – dark (active) phase; yellow – light (rest) phase. For ECoG PSD figures, data represent mean percentage change in ECoG PSD (0.1-25 Hz) ± S.E.M. Statistical significance is indicated by the following symbols: * – dose main effect; # – dose x time interaction effect; † – linear trend contrast; ^ pairwise comparison; × – 0.1 mg·kg^-1^ oxytocin vs VEH; + – 0.3 mg·kg^-1^ oxytocin vs VEH; ∼ – 1 mg·kg^-1^ oxytocin vs VEH; • – 10 mg·kg^-1^ caffeine vs VEH. For graphs with two y-axes, the alignment of the significance symbol represents which outcome the significance refers to: left – bout frequency; right – mean bout duration. Level of statistical significance is indicated by the number of symbols: one – *p* < .05, two – *p* < .01, three – *p* < .001, four – *p* < .0001.

Overall, caffeine (i.p.; 10 mg·kg^-1^) reduced %NREMS, and this differed over time during the recording session (Figure 3Bi). During 0-90 min, caffeine greatly reduced %NREMS (Figure 3Av), due to both reduced NREMS bouts and reduced mean NREMS bout duration (Figure 3Avi). Additionally, caffeine increased sleep onset latency (Figure 3Avii).

During the 1^st^ hour post-dose, relative to vehicle, 1 mg·kg^-1^ oxytocin reduced ECoG PSD in the theta, alpha, sigma, and beta frequency bands, while caffeine reduced ECoG PSD across all frequency bands (Figure 3Bii). In contrast, during the last 5 hours of recording, oxytocin increased ECoG PSD in the delta (all doses), theta (all doses), alpha (all doses), and beta (0.1 mg·kg^-1^) frequency bands, whereas caffeine had no effect on ECoG PSD (Figure 3Biii).

##### REM sleep

Overall, oxytocin influenced %REMS, and this differed over time during the recording session (Figure 3Ci). During 30-180 min, oxytocin dose-dependently reduced %REMS (Figure 3Cii), due to reductions in both REMS bouts and mean REMS bout duration (Figure 3Ciii). Additionally, oxytocin dose-dependently increased REMS onset latency (Figure 3Civ).

Overall, caffeine reduced %REMS, and this effect varied over the course of the recording (Figure 3Di). During 30-180 min, caffeine reduced %REMS (Figure 3Cv), due to both reduced REMS bouts and mean REMS bout duration of REM bouts (Figure 3Cvi). Furthermore, caffeine increased REMS onset latency (Figure 3Cvii).

Over the 6-hour recording, relative to vehicle, oxytocin increased ECoG PSD in the delta (0.1 and 1 mg·kg^-1^), alpha (all doses), sigma (0.1 and 1 mg·kg^-1^), and beta (all doses) frequency bands. In contrast, caffeine reduced ECoG PSD in the sigma and beta frequency bands (Figure 3Dii).

#### Body temperature

Administration of i.p. oxytocin dose-dependently reduced body temperature (Figure S1A and B). In contrast, i.p. caffeine (10 mg·kg^-1^) slightly elevated body temperature (Figure S1C and D).

### Experiment 2 – Influence of oxytocin receptor antagonism on oxytocin-induced effects on sleep-wake outcomes

No differences were observed in the proportion of rats in oestrus phases between dose conditions and their respective controls (Table S8). Summarised statistical results for Experiment 2 are presented in Table 2 and S7.

#### Active wake

From 0-30 min, pre-administration of the selective oxytocin receptor antagonist L-368,899 (i.p.; 5 mg·kg^-1^) inhibited the oxytocin-induced (i.p.; 1 mg·kg^-1^) increase in bouts of AW (Figure 4Aii) but not in %AW (Figure 4Ai) or mean duration of AW bouts (Figure 4Aiii). From 30-180 min, pre-administration of L-368,899 significantly attenuated the oxytocin-induced increase in %AW and bouts of AW (Figures 4Aiv and Av) but not mean duration of AW bouts (Figure 4Avi).

**Figure 4.**
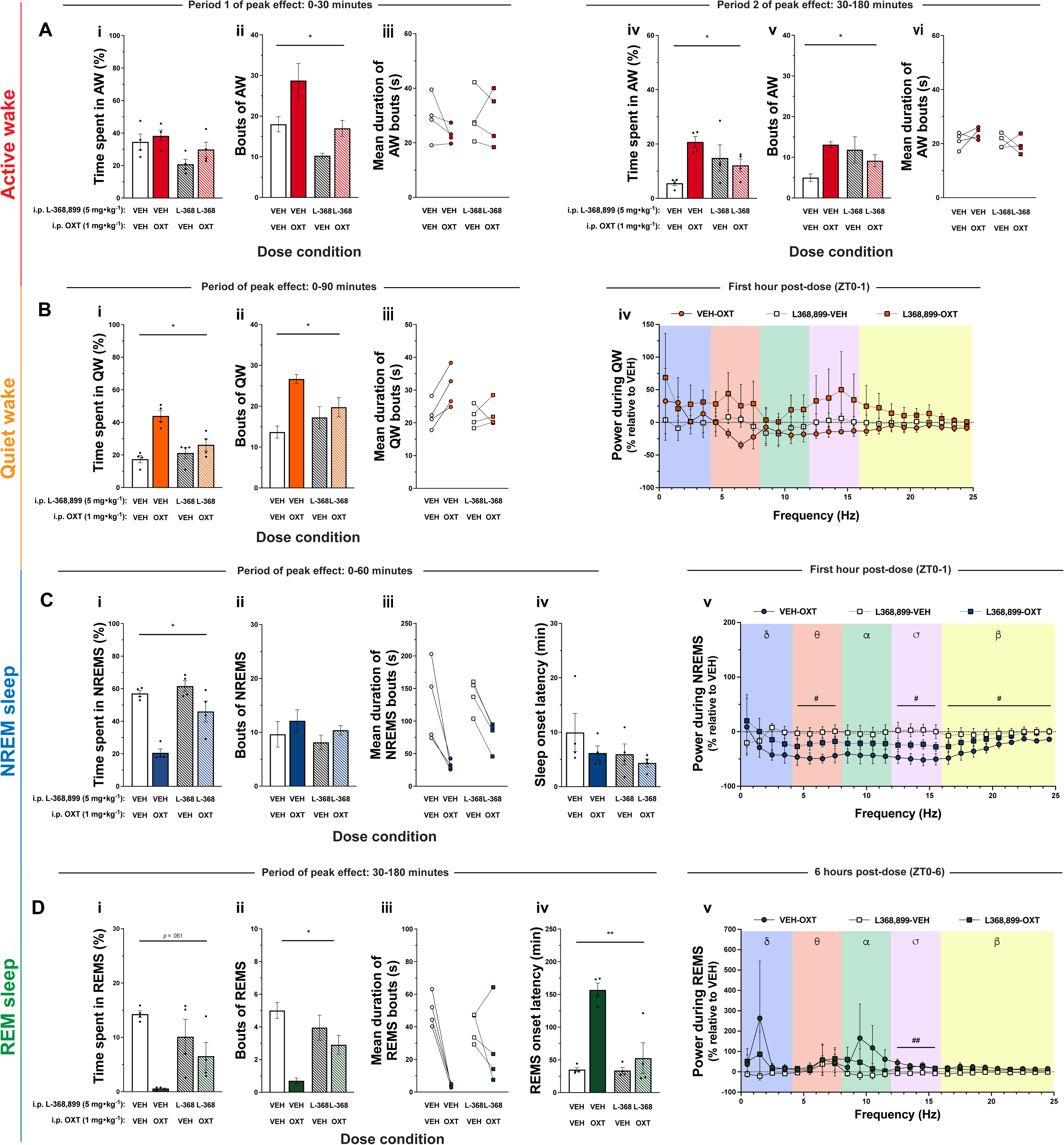
Influence of OXTR antagonism on oxytocin-induced effects on sleep-wake outcomes. Influence of pre-administration of the OXTR antagonist L-368,899 (i.p.; 5 mg·kg^-1^) on oxytocin-induced (i.p.; 1 mg·kg^-1^) effects on: %AW **(Ai)**, AW bouts **(Aii)**, and mean duration of AW bouts **(Aiii)** during first period of peak effect (0-30 min); %AW **(Aiv)**, AW bouts **(Av)**, and mean duration of AW bouts **(Avi)** during second period of peak effect (30-180 min); %QW **(Bi)**, QW bouts **(Bii)**, and mean duration of QW bouts **(Biii)** during the period of peak effects (0-90 min); ECoG power spectral density from 0-1 h during QW **(Biv)**; %NREMS **(Ci)**, NREMS bouts **(Cii)**, and mean duration of NREMS bouts **(Ciii)** during the period of peak effects (0-60 min); sleep onset latency **(Civ)**; ECoG power spectral density from 0-1 h during NREMS **(Cv)**; %REM sleep **(Di)**, REMS bouts **(Dii)**, and mean duration of REMS bouts **(Diii)** during period of peak effect (30-180 min); REMS onset latency **(Div)**; and ECoG power spectral density from 0-6 h during REMS **(Dv)**. Sample size for all dose conditions is *n* = 4 females. Data represent mean values ± S.E.M., individual data points represent individual subject data (closed circles – females). For **Aiii**, **Avi**, **Biii**, **Ciii**, and **Diii**, data points represent individual subject data with lines connecting same subject data. Antagonist was administered at ZT23.75 and oxytocin was administered at ZT0. For ECoG PSD figures, data represent mean percentage change in ECoG PSD (0.1-25 Hz) ± S.E.M. Data represent mean percentage change in ECoG PSD (0-25 Hz) ± S.E.M. Statistical significance is indicated by the following symbols: * – dose x antagonist interaction effect; # – oxytocin-VEH vs VEH-VEH (baseline) comparison. For graphs with two y-axes, the alignment of the significance symbol represents which outcome the significance refers to: left – bout frequency; right – mean bout duration. Level of statistical significance is indicated by the number of symbols: one – *p* < .05, two – *p* < .01, three – *p* < .001, four – *p* < .0001.

#### Quiet wake

Pre-administration of L-368,899 prevented the oxytocin-induced increase in %QW (Figure 4Bi) and QW bout frequency (Figure 4Bii), but did not reach significance for mean QW bout duration (Figure 4Biii). No significant effects of any condition on ECoG PSD during were detected at any frequency band (Figure 4Biv).

#### NREM sleep

Pre-administration of L-368,899 antagonised the oxytocin-induced reduction in %NREMS (Figure 4Ci); effects of antagonism on NREMS bout frequency, mean duration of NREMS bouts, and sleep onset latency were not clear (Figures 4Cii, Ciii, and Civ). Relative to vehicle, oxytocin reduced ECoG PSD within the theta, sigma, and beta frequency bands, and these changes were not observed when L-368,899 was pre-administered before oxytocin (Figure 4Cv).

#### REM sleep

Pre-administration of L-368,899 antagonised the oxytocin-induced reduction in REMS bout frequency (Figure 4Dii) and REMS onset latency (Figure 4Div), but did not reach significance for %REM sleep (Figure 4Di, *p* = 0.051) or mean REMS bout duration (Figure 4Diii). Relative to vehicle, while oxytocin significantly increased ECoG PSD within the sigma frequency band, no significant effect of co-administration of oxytocin-L-368,899 or L-368,899 alone on ECoG PSD was found for any frequency band (Figure 4Dv).

#### Body temperature

Administration of i.p. oxytocin (1 mg·kg^-1^) reduced body temperature (Figure S2). Notably, pre-administration of L-368,899 attenuated this reduction in body temperature (see Supplemental material for details).

### Experiment 3 – Effects of i.n. oxytocin and i.n. caffeine on sleep-wake outcomes

There were no differences in the proportion of rats in different oestrus phases between oxytocin dose conditions and VEH. However, the proportion of rats in different oestrus phases significantly differed between caffeine and VEH (see Tables S10 and S11). No main effects of dose across %sleep-wake state outcomes at baseline were observed (see Table S12 and S13). Summarised statistical results for Experiment 3a and 3b are presented in Tables 3 and S9, respectively.

#### Wake outcomes

*Active wake.* There were no significant effects of i.n. oxytocin (at any dose) on any AW outcome (Figure 5Ai-v and Bi-v). Averaged across the 6-h recording, caffeine (i.n., 10 mg·kg^-1^) increased %AW (Figure 5Bi). During 0-30 min, caffeine increased %AW (Figure 5Aiv), but effects on AW bouts and mean AW bout duration were not statistically significant (Figure 5Av). From 30-180 min, caffeine increased %AW (Figure 5Biv) and AW bouts (Figure 5Bv), but did not increased mean AW bout duration.

**Figure 5.**
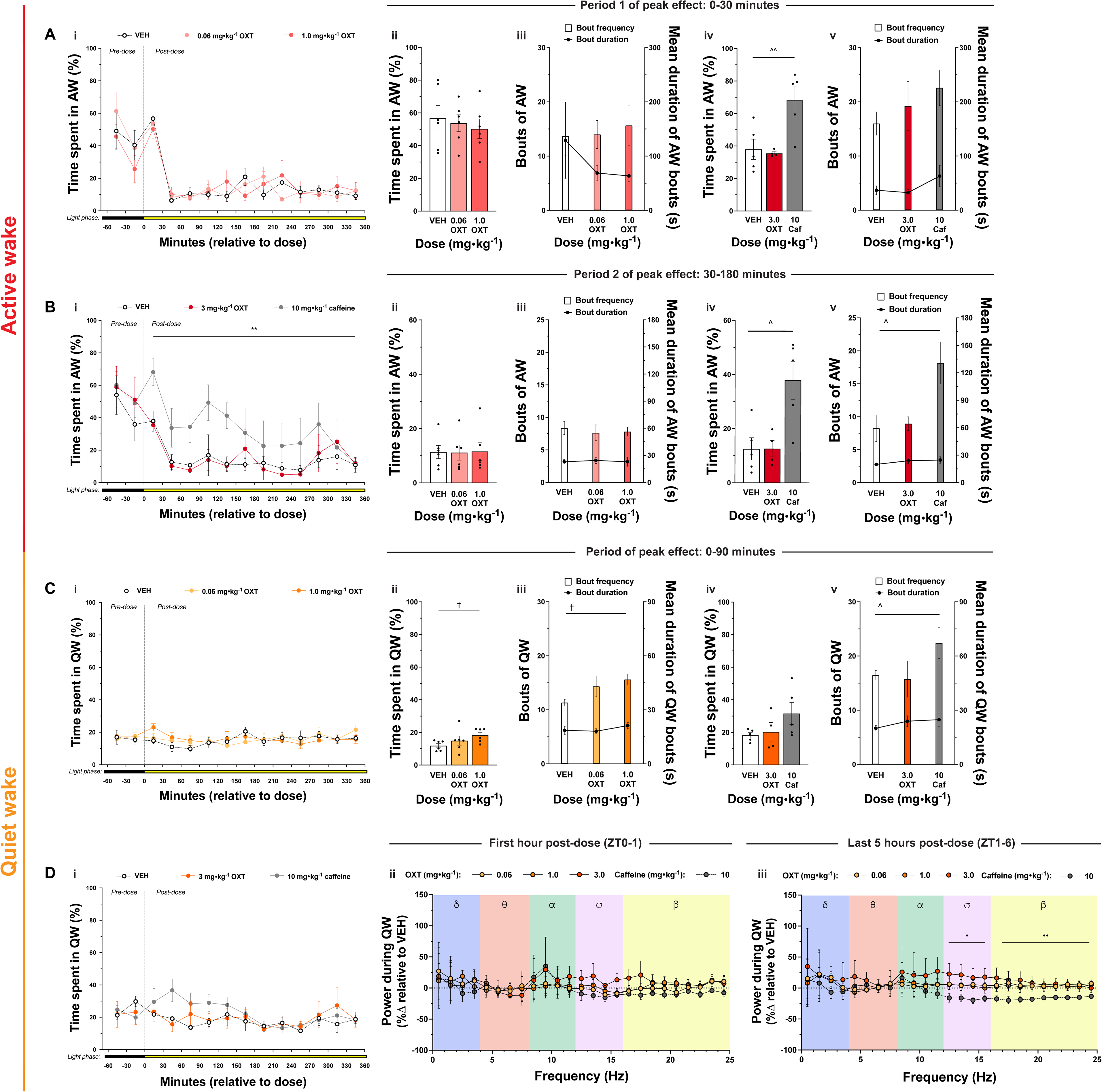
Effects of i.n. oxytocin and i.n. caffeine on wake outcomes. *Low and mid dose oxytocin effects.* Influence of i.n. oxytocin (0, 0.06, and 1 mg·kg^-1^) on %AW across entire 7-hour recording session **(Ai)**, %AW during first period of peak effect (0-30 min) **(Aii)**, AW bout frequency and mean AW bout duration during first period of peak effect (0-30 min) **(Aiii)**, %AW during second period of peak effect (30-180 min) **(Bii)**, AW bout frequency and mean AW bout duration during second period of peak effect (30-180 min) **(Biii)**, %QW across entire 7-hour recording session **(Ci)**, %QW during first period of peak effect (0-90 min) **(Cii)**, and QW bout frequency and mean QW bout duration during first period of peak effect (0-90 min) **(Ciii)**. *High dose oxytocin and caffeine effects.* Influence of i.n. oxytocin (3 mg·kg^-1^) and i.n. caffeine (10 mg·kg^-1^) on %AW across entire 7-hour recording session **(Bi)**, %AW during first period of peak effect (0-30 min) **(Aiv)**, AW bout frequency and mean AW bout duration during first period of peak effect (0-30 min) **(Av)**, %AW during second period of peak effect (30-180 min) **(Biv)**, AW bout frequency and mean AW bout duration during second period of peak effect (30-180 min) **(Bv)**, %QW across entire 7-hour recording session **(Di)**, %QW during period of peak effect (0-90 min) **(Civ)**, and QW bout frequency and mean QW bout duration during period of peak effect (0-90 min) **(Cv)**. *Power spectral density effects.* Influence of i.n. oxytocin and caffeine on ECoG power spectral density from 0-1 h **(Dii)** and 1-6 h **(Diii)** during QW. Sample size for Experiment 3a was *n* = 6 females for all doses. Sample size for Experiment 3b was *n* = 5 females (for all doses) except for the 3 mg·kg^-1^ oxytocin dose (*n* = 4). Data represent mean values ± S.E.M., individual data points represent individual subject data (closed circles – females). For % time spent in sleep-wake state graphs, data points represent values for 30-min bins; values pertain to the bin of time defined by the x-axis timepoints to the immediate left and right of the data point. Dose was administered at ZT0 and colour of bar below x-axis signifies light cycle phase at each time point: black – dark (active) phase; yellow – light (rest) phase. For ECoG PSD figures, data represent mean percentage change in ECoG PSD (0.1-25 Hz) ± S.E.M. Data represent mean percentage change in ECoG PSD (0-25 Hz) ± S.E.M. Statistical significance is indicated by the following symbols: * – dose main effect; † – linear trend contrast; ^ pairwise comparison; • – 10 mg·kg^-1^ caffeine vs VEH. For graphs with two y-axes, the alignment of the significance symbol represents which outcome the significance refers to: left – bout frequency; right – mean bout duration. Level of statistical significance is indicated by the number of symbols: one – *p* < .05, two – *p* < .01, three – *p* < .001, four – *p* < .0001.

#### Quiet wake

Averaged across the entire recording, there was no significant effect of oxytocin at any dose on %QW (Figure 5Ci and Di). From 0-90 min, oxytocin significantly—albeit slightly—increased %QW in a dose-dependent manner across 0, 0.06 and 1 mg·kg^-1^ doses (Figure 5Cii), due to dose-dependent increases in QW bouts rather than QW bout duration (Figure 5Ciii). However, this increase in %QW and QW bouts was not found for the higher 3 mg·kg^-1^ oxytocin dose (Figure 5Civ and Cv). Additionally, no effects of oxytocin on ECoG PSD during QW were found at any dose at any time (Figure 5Dii and Diii).

Averaged across the entire session, no significant effect of caffeine on %QW was observed (Figure 5Di). However, from 0-90 min, caffeine increased the number of QW bouts but not %QW or mean QW bout duration (Figures 5Civ and Cv). During the last 5 hours of recording, i.n. caffeine significantly reduced ECoG PSD relative to vehicle in the sigma and beta frequency bands (Figure 5Diii), however, no other significant effects of caffeine on QW PSD outcomes were observed.

#### Sleep outcomes

*NREM sleep.* No significant effects of i.n. oxytocin (at any dose) on any NREMS outcomes were found (Figure 6Ai-vii and Bi-iii). Averaged across the 6-hour session, i.n. caffeine (10 mg·kg^-1^) significantly reduced %NREMS (Figure 6Bi); the increase in sleep onset latency did not reach statistical significance (Figure 6Avii). During 0-90 min, i.n. caffeine reduced %NREMS (Figure 6Av) and reduced duration of NREMS bouts (Figure 6Avi), but did not significantly reduce bout frequency. During the first hour of recording, relative to vehicle, caffeine significantly reduced ECoG PSD within the delta, theta, alpha and sigma frequency bands (Figure 6Bi), however no effect of caffeine was detected over the following 5 hours of recording (Figure 6Bii).

**Figure 6.**
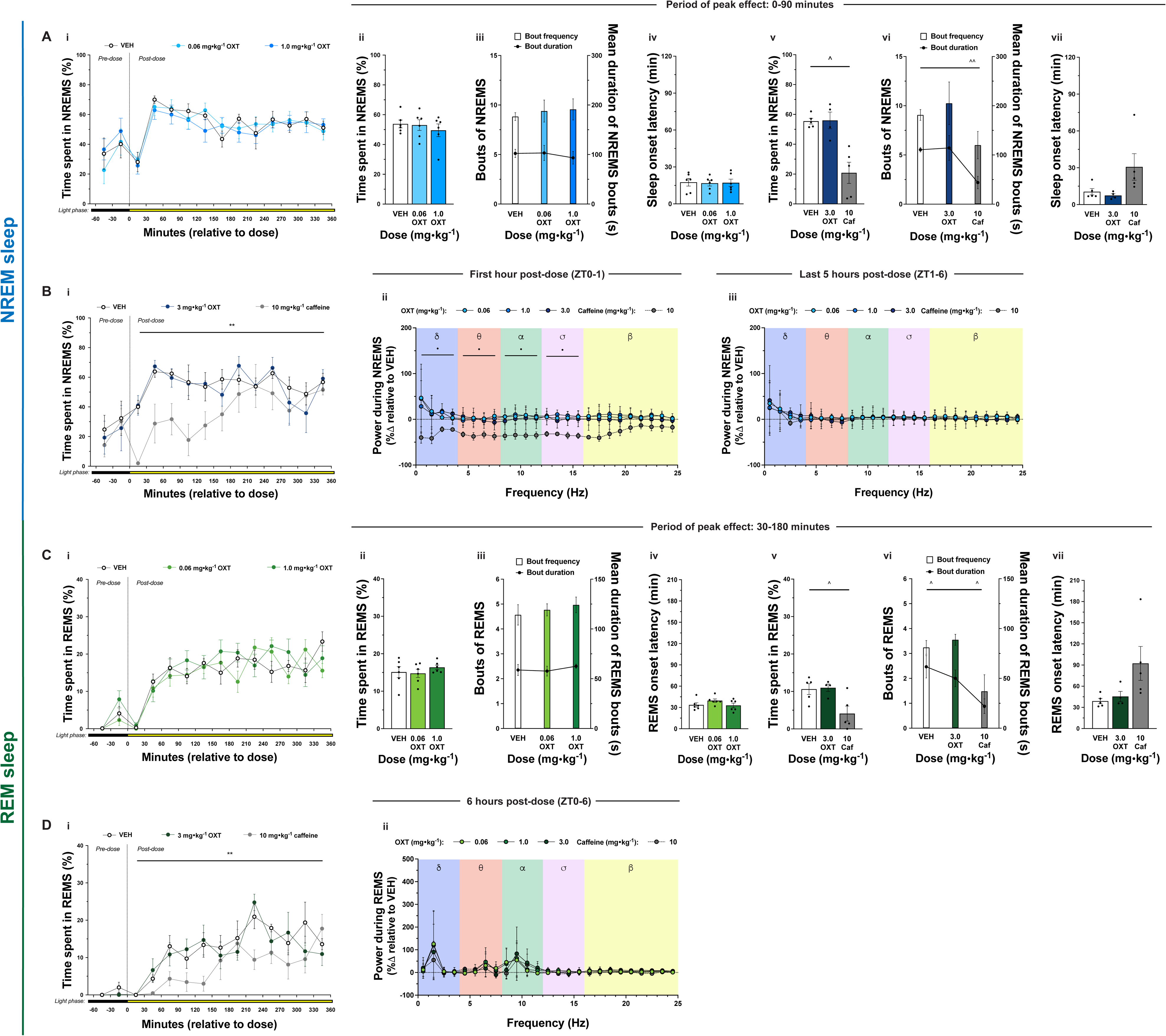
Effects of i.n. oxytocin and i.n. caffeine on sleep outcomes. *Lower oxytocin dose effects.* Influence of i.n. oxytocin dose range (0, 0.06, 1 mg·kg^-1^) on %NREMS across entire 7-hour recording session **(Ai)**, %NREM during period of peak effect (0-90 min) **(Aii)**, NREMS bout frequency and mean NREMS bout duration during period of peak effect (0-90 min) **(Aiii)**, sleep onset latency **(Aiv)**, %REM sleep across entire 7-hour recording session **(Ci)**, %REM sleep during period of peak effect (30-180 min) **(Cii)**, and REMS bout frequency and mean REMS bout duration during period of peak effect (30-180 min) **(Ciii)**, and REMS onset latency **(Civ)**. *Higher oxytocin dose and caffeine effects.* Influence of i.n. oxytocin (3 mg·kg^-1^) and i.n. caffeine (10 mg·kg^-1^) on %NREMS across entire 7-hour recording session **(Bi)**, %NREMS during period of peak effect (0-90 min) **(Av)**, NREMS bout frequency and mean NREMS bout duration during period of peak effect (0-90 min) **(Aii)**, sleep onset latency **(Cvii)**, %REM sleep across entire 7-hour recording session **(Di)**, %REM sleep during period of peak effect (30-180 min) **(Cv)**, REMS bout frequency and mean REMS bout duration during period of peak effect (30-180 min) **(Cvi)**, and REMS onset latency **(Cvii)**. *Power spectral density effects.* Influence of i.n. oxytocin dose range and caffeine on ECoG power spectral density from 0-1 h **(Bii)** and 1-6 h **(Biii)** during NREMS and 0-6 h during REMS **(Dii)**. Sample size for Experiment 3a was *n* = 6 females for all doses. Sample size for Experiment 3b was *n* = 5 females (for all doses) except for 3 mg·kg^-1^ oxytocin which was *n* = 4. Data represent mean values ± S.E.M., individual data points represent individual subject data (closed circles – females). For % time spent in sleep-wake state graphs, data points represent values for 30-min bins; values pertain to the bin of time defined by the x-axis timepoints to the immediate left and right of the data point. Dose was administered at ZT0 and colour of bar below x-axis signifies light cycle phase at each time point: black – dark (active) phase; yellow – light (rest) phase. For ECoG PSD figures, data represent mean percentage change in ECoG PSD (0.1-25 Hz) ± S.E.M. Data represent mean percentage change in ECoG PSD (0-25 Hz) ± S.E.M. Statistical significance is indicated by the following symbols: * – dose main effect; ^ pairwise comparison; • – 10 mg·kg^-1^ caffeine vs VEH. For graphs with two y-axes, the alignment of the significance symbol represents which outcome the significance refers to: left – bout frequency; right – mean bout duration. Level of statistical significance is indicated by the number of symbols: one – *p* < .05, two – *p* < .01, three – *p* < .001, four – *p* < .0001.

#### REM sleep

No significant effects of i.n. oxytocin (at any dose) on any REMS outcomes were found (Figure 6Ci-vii and 6Di-ii). Overall, i.n. caffeine (10 mg·kg^-1^) significantly reduced %REM sleep (Figure 6Di), however the increase in REMS onset latency did not reach statistical significance (Figure 6Cvii). During 30-180 min, i.n. caffeine reduced %REM sleep (Figure 6Cv), due to both reduced REMS bout frequency and mean REMS bout duration (Figure 6Cvi). Relative to vehicle, no effect of caffeine on ECoG PSD across recording sessions was detected (Figure 6Dii).

#### Body temperature

Administration of i.n. oxytocin did not significantly affect body temperature at any dose. Likewise, no significant effect of i.n. caffeine on body temperature was found (Figure S3).

## Discussion

The current study explored the acute effects of exogenous oxytocin and caffeine, administered via two routes, on sleep-wake outcomes, and whether observed effects were mediated by the OXTR. As hypothesised, i.p. oxytocin promoted wakefulness and suppressed sleep in a *dose-dependent* manner in both female and male rats. I.p. oxytocin predominantly promoted a state of quiet wakefulness, reduced AW, suppressed NREMS and REMS, and delayed REMS onset. Relative to oxytocin, caffeine effects on AW and NREMS appeared more pronounced, on QW less pronounced, and on REMS of similar magnitude. I.p. oxytocin-induced sleep-wake effects were at least partially mediated by the OXTR. I.n. oxytocin failed to recapitulate most i.p. oxytocin-induced effects on sleep-wake behaviour across the dose range tested, whereas i.n. and i.p. caffeine had similar effects. Oxytocin-induced sleep-wake effects therefore appear dependent on dose and route of administration, but not biological sex.

### Timing and dose-dependency

I.p. oxytocin had clear dose-dependent effects on almost all sleep-wake outcomes. Immediately post-administration, oxytocin facilitated an exchange of AW for QW, without impacting sleep; in contrast, caffeine increased AW at the cost of NREMS. From 30-90 min, oxytocin maintained substantial elevations in QW, primarily at the cost of NREMS and REMS sleep. During QW and NREMS within this wake-dominant period, both caffeine and oxytocin (1 mg·kg^-1^) reduced ECoG PSD across frequency bands reflecting reduced homeostatic sleep pressure ^26, 43, 44^. This oxytocin-induced wake-promotion is unlikely due to disruptions in thermoregulation; while oxytocin reduced body temperature, this is typically associated with *improved* NREMS induction^45^.

Although oxytocin-induced NREMS suppression abated by approximately 90 min post-administration with no observed rebound sleep, ECoG PSD in lower frequency bands (delta, theta, alpha) remained elevated for the last 5 hours of recording. This aligns with Lancel, Krömer and Neumann ^20^ who found similar effects of i.c.v. oxytocin on EEG PSD. Notably, this elevation in lower frequency PSD occurred for all i.p. oxytocin doses tested and likely reflects increased sleep pressure ^46^, increased sleep intensity ^39^, or sleep propensity ^43^. While a brief elevation following oxytocin-induced NREMS suppression might suggest a transient rebound in sleep pressure, the protracted nature of this effect is unlikely due to homeostatic sleep drive. Tobler and Borbély ^47^ demonstrated that 3-h total sleep deprivation in rats—a relatively longer, more potent sleep deprivation than the current study—had no significant impact on EEG PSD within the delta frequency (0.75-4.5 Hz) band during NREMS. Hence, oxytocin may induce persisting increases in sleep intensity during NREMS, independent of initial NREMS suppression. Caffeine reduced sleep pressure during NREMS, as expected ^26^.

The most prolonged behavioural effect of i.p. oxytocin was the suppression of REMS. Since NREMS typically precedes REMS during normal sleep architecture ^37^, oxytocin-induced disruption to NREMS would be expected to impact REMS. However, while caffeine delayed both sleep- and REMS-onset latency, and effects on NREMS and REMS were mostly temporally co-occurring, oxytocin only impacted REMS-onset latency, and REMS effects endured considerably beyond NREMS disruptions. This suggests that oxytocin dose-dependently suppresses transitions from NREMS to REMS via mechanisms independent from simply disrupting natural progression through the sleep cycle.

### Quiet wakefulness and mechanisms

Oxytocin-induced promotion of quiet wakefulness incorporates aspects of both competing hypotheses that motivated this study: *enhanced environmental awareness* (wake-promotion) and *stress attenuation-quiescence* (sleep-promotion). The observed quiet wakefulness was a state of sustained arousal, minimal locomotion, reduced physical activity, rest, and relaxation. This result aligns with Lancel, Krömer and Neumann ^20^, who found central administration of oxytocin in rats promoted wakefulness at the cost of sleep. However, that study did not delineate between active and quiet wakefulness. The current findings may also align with research by Mahalati, Okanoya, Witt and Carter ^22^, who found i.c.v. oxytocin promoted sleep-like postures in prairie voles. While this was interpreted as oxytocin-induced sleep-promotion, the lack of polysomnographic assessment and reliance on postural observation would have rendered quiet wakefulness difficult to distinguish from NREMS.

Within the preclinical literature concerning the quiet wake state, various titles are used (i.e., resting wake, awake quiescence, attentive wake) and defining criteria are typically vague, differing within and between species. However, most definitions converge on the fundamental criterion of ‘relaxed’ wakefulness with no locomotion ^48–51^. Recently, studies using PSG-based definitions demonstrated important neurobiological differences between active and quiet wakefulness: beta activity (15-35 Hz)—during QW but not AW—couples with slow-wave activity to reveal homeostatic sleep drive ^52^.

Furthermore, Moriya, Kanamaru, Okuma et al.^51^ showed that both QW and NREM sleep share similar patterns of acetylcholine and glutamate signalling in the hippocampus, and that experimental interventions can independently impact active wakefulness without influencing quiet wakefulness.

Previous studies have implicated the pontine parabrachial nucleus (PBn) in quiet wakefulness. Qiu, Chen, Fuller and Lu ^53^ demonstrated that chemogenetic activation of parabrachial neurons elicited a “behaviourally quiet” wakefulness and “alert but minimally active arousal” at the cost of NREMS and REMS during the light phase. This wake-promoting effect was mediated via parabrachial projections to the lateral hypothalamus and basal forebrain. Notably, although these authors do not use the term quiet wake, when compared to spontaneous wake, representative traces of the “quiet” state indicate lower EMG amplitude and more regular EMG tone ^53^, consistent with our QW definition in the current study. More recently, Xu, Wang, Dong et al.^54^ found that chemogenetic and optogenetic activation of medial parabrachial neurons in rats induced an attentive wake state (i.e., wake with no locomotion, some head movement, and low EMG tone) via glutamatergic projections to the LH and BF. Since OXTRs are expressed by PBn neurons and as the PBn receives direct projections from PVN oxytocinergic neurons ^55^, it is possible that exogenous oxytocin induces quiet wakefulness through PBn-mediated mechanisms. While unspecific to *quiet* wakefulness, several other potential mechanisms for oxytocin-induced wakefulness exist. These include an oxytocin-orexin positive feedback interaction within the LH ^21^ and co-transmission of glutamate and corticotropin-releasing hormone (CRH) from oxytocinergic neurons ^2^.

The OXTR antagonist L-368,899 blocked some, but not all sleep-wake effects. It is possible that a higher dose of the antagonist was needed for complete blockade. However, given oxytocin demonstrates considerable affinity for the V1aR ^56^, some effects of exogenous oxytocin are V1aR-mediated ^15, 19, 57, 58^, and the vasopressin system is also involved in regulation of sleep-wake behaviours ^59, 60^, future studies might also explore potential co-contribution of OXTR-V1aR activation in oxytocin sleep-wake effects.

It is also important to consider the potential for peripheral mediation of oxytocin-induced wakefulness effects. Although peripheral oxytocin administration can increase central oxytocin levels ^33, 61^, peripheral oxytocin also exerts some effects through stimulation of the ascending vagal afferent pathway ^62–64^ and vagal stimulation can promote wakefulness ^65–67^. However, given previous work ^20^ found i.c.v. oxytocin also promotes wakefulness, central mediation of at least some of the observed effects of i.p. oxytocin in the present study seems likely. Nevertheless, future research might investigate the relative contribution of central and peripheral targets in oxytocin sleep-wake effects.

### Oxytocin and route of administration – intraperitoneal vs intranasal

Exogenous oxytocin-induced sleep-wake effects were dependent on route of administration. At equivalent or higher doses, i.n. oxytocin failed to replicate i.p. oxytocin effects on AW, NREMS and REMS outcomes, and only exerted a relatively small increase in QW compared to the large effect induced by i.p. oxytocin. Since equivalent doses of i.p. and i.n. caffeine exerted comparable effects, this route of administration effect should not be interpreted as an issue with the i.n. administration procedure. Rather, this finding may be due to poorer bioavailability and/or differential distribution of oxytocin to sleep-relevant central or peripheral targets with i.n. oxytocin administration ^68^.

There is equivocal evidence regarding both the differential bioavailability and distribution of i.n. administered oxytocin, compared to other routes. Compared to oxytocin administered via i.p. and/or intravenous (i.v.) routes, i.n. oxytocin administered at the same dose has been found to produce lower 1.5-fold lower^33^, 20-fold lower^69^ or comparable ^61^ plasma oxytocin levels in rodents. Higher oxytocin levels tend to be observed in olfactory regions following i.n. administration ^69, 70^ and one study reported higher oxytocin levels in the amygdala and hippocampus with i.p. compared to i.n. oxytocin ^33^, although this was not observed in another study ^61^.

Hence, it is possible that in the current study, i.p. oxytocin reached functionally relevant concentrations in sleep-wake regulatory brain regions or peripheral targets and i.n. oxytocin did not. The comparable effects of i.n. and i.p. caffeine suggest similar distribution via both routes of administration and possible fundamental differences in pharmacokinetics between caffeine and oxytocin ^71^. It could simply be that substantially higher doses of i.n. oxytocin are required. Clearly, more research is needed examining the differential bioavailability and distribution of oxytocin with different routes of administration.

The current study possessed several strengths in design and methodology: (1) the inclusion of caffeine as a wake-promoting positive control facilitated crucial informative comparisons to oxytocin, (2) both male and female rats were tested ^72^, (3) co-administration of oxytocin with a centrally penetrant OXTR antagonist facilitated determination of OXTR involvement, (4) i.n. administration was conducted without anaesthetic to avoid known interference with sleep and circadian rhythms ^73^, and (5) wireless telemetry was used for PSG recording to avoid potential interference of tethering and restriction of movement with sleep-wake behaviour ^74^.

Understanding the impact of oxytocin—both endogenous and exogenous—on sleep-wake behaviour holds valuable implications for basic and applied research. Our research confirms exogenous oxytocin can induce wakefulness and that mixed preclinical evidence on oxytocin effects on sleep-wake outcomes is likely due to inter-study differences in dose and route of administration ^2^. Given growing interest in the development of oxytocin system targeting pharmacotherapeutics ^75^ and use of i.n. oxytocin in clinical trials for psychiatric disorders ^76^, this study also offers novel insight into potential therapeutic applications. Inducing quiet wakefulness could prove valuable for disorders involving agitation and aggression (e.g., dementia)^77^ and excessive daytime sleepiness (e.g., idiopathic hypersomnolence, OSA, Prader-Willi Syndrome, and narcolepsy)^78^.

More broadly, whilst the present study focused on the effects of exogenous oxytocin, it highlights the potential for the endogenous oxytocin system to regulate sleep-wake behaviours. Mammalian sleep mostly occurs within a social context (i.e., co-sleeping)^79^, which often includes affiliative behaviours that involve endogenous oxytocin signalling^80, 81^. Thus, understanding the role of endogenous oxytocin in sleep-wake behaviour may elucidate how co-sleeping impacts sleep quality, both objectively and subjectively ^79^, and how alterations to endogenous oxytocin may impact sleep.

The current study reconciles the apparent contradictory *environmental awareness-arousal* (wake-promotion) and *stress attenuation-quiescence* (sleep-promotion) hypotheses of oxytocin-induced influences on sleep-wake behaviour. Exogenous oxytocin acutely promotes quiet wakefulness, a state of restfulness and sustained arousal to support environmental awareness, at the cost of active wakefulness, NREMS and REMS. Importantly, these effects are dependent on dose and route of administration, but not biological sex.

## Data availability statement

The data that support the findings of this study are available from the corresponding author upon reasonable request.

## Author contributions

Author contributions, as defined by the Contributor Roles Taxonomy (CRediT) matrix, are detailed as follows: Conceptualisation (JSR and MTB), ethics application (JSR, NAE, AG, and MTB), data curation (JSR and NAE), formal analysis (JSR, AG, and MTB), funding acquisition (MTB), investigation (JSR and NAE), methodology (JSR, NAE, AG, and MTB), project administration (JSR and MTB), resources (NAE, AG, MTB), software (JSR), supervision (NAE, AG, MTB), validation (JSR and MTB), visualisation (JSR and MTB), writing – original draft (JSR and MTB), and writing – review and editing (JSR, NAE, AG, and MTB).

## Ethics statement

The experiments reported in this study were conducted in line with the guidelines laid out in the *Australian code for the care and use of animals for scientific purposes (8^th^ edition, 2013)* and were approved by the Animal Ethics Committee at The University of Sydney (AEC number: 2019/1615). All experiments are reported in compliance with guidelines of ARRIVE 2.0 guidelines those specified by the British Journal of Pharmacology.

## Acknowledgements

We are grateful to Erin Lynch and Cassandra Hanbury-Brown for their insight into interpretation of ECoG data, and to technical staff Stephan Martin and Yann Abéguilé at Data Science International for their advice on conducting sleep scoring and analysis of ECoG data. Additionally, we would like to thank Jennifer McKenna, Vincent Zappala and Laboratory Animal Services administrative staff at The University of Sydney for their assistance during the study and provision of animal welfare.

