## Supplementary material for "Quiet wakefulness: The influence of intraperitoneal and intranasal oxytocin on sleep-wake behaviour and neurophysiology in rats": Tables 1-3

**Table 1 – Effects of i.p. oxytocin dose range and i.p. caffeine dose on sleep-wake outcomes.**

| Wake outcome | Oxytocin dose-response (0, 0.1, 0.3, 1 mg·kg$^{-1}$) | | | Caffeine 10 mg·kg$^{-1}$ (positive control) | |
|---|---|---|---|---|---|
| | ***Linear trend*** | ***Main effect of dose*** | ***Dose x Time interaction*** | ***Main effect of dose*** | ***Dose x Time interaction*** |
| ***Active wake*** | | | | | |
| Proportion of time | $F(1, 38) = 0.87$ $p = .3570$ | $F(2.302, 29.93) = 0.55$ $p = .6070$ | $F(8.546, 108.0) = 3.85$ $p = .0004$ | $F(1, 11) = 68.56$ $p < .0001$ | $F(5.006, 55.06) = 6.79$ $p < .0001$ |
| Bout frequency | $F(1, 38) = 13.90$ $p = .0006$ | $F(2.847, 37.01) = 4.13$ $p = .0139$ | $F(7.608, 96.13) = 3.57$ $p = .0014$ | $F(1, 11) = 67.86$ $p < .0001$ | $F(5.079, 55.86) = 3.38$ $p = .0095$ |
| Bout duration | $F(1, 38) = 3.00$ $p = .0911$ | $F(1.891, 24.58) = 0.89$ $p = .4189$ | $F(3.016, 38.11) = 2.76$ $p = .0549$ | $F(1, 11) = 0.88$ $p = .3679$ | $F(3.018, 33.20) = 2.59$ $p = .0692$ |
| *AW 0-30 min* | | | | | |
| Proportion of time | $F(1, 38) = 43.46$ $p < .0001$ | $F(2.116, 26.80) = 15.37$ $p < .0001$ | – | $t(11) = 5.20$ $p = .0003$ | – |
| Bout frequency | $F(1, 38) = 15.64$ $p = .0003$ | $F(2.437, 30.87) = 6.52$ $p = .0027$ | – | $t(11) = 1.71$ $p = .1148$ | – |
| Bout duration | $F(1, 38) = 15.49$ p = .0003 | $F(1.259, 15.95) = 5.56$ $p = .0253$ | – | $t(11) = 1.49$ $p = .1653$ | – |
| *AW 30-180 min* | | | | | |
| Proportion of time | $F(1, 38) = 44.09$ $p < .0001$ | $F(2.313, 29.30) = 15.25$ $p < .0001$ | – | $t(11) = 7.45$ $p < .0001$ | – |
| Bout frequency | $F(1, 38) = 35.75$ $p < .0001$ | $F(2.513, 31.83) = 12.86$ $p < .0001$ | – | $t(11) = 9.96$ $p < .0001$ | – |
| Bout duration | $F(1, 38) = 5.83$ $p = .0207$ | $F(1.836, 23.26) = 2.41$ $p = .1154$ | – | $t(11) = 1.40$ $p = .1892$ | – |
| ***Quiet wake*** | | | | | |
| Proportion of time | $F(1, 38) = 22.23$ $p < .0001$ | $F(2.451, 31.87) = 7.55$ $p = .0011$ | $F(7.515, 94.96) = 8.06$ $p < .0001$ | $F(1, 11) = 14.31$ $p = .0030$ | $F(4.655, 51.21) = 2.79$ $p = .0293$ |
| Bout frequency | $F(1, 38) = 12.11$ $p = .0013$ | $F(2.372, 30.84) = 1.98$ $p = .1491$ | $F(8.618, 108.9) = 2.41$ $p = .0172$ | $F(1, 11) = 33.24$ $p = .0001$ | $F(5.113, 56.24) = 2.67$ $p = .0303$ |
| Bout duration | $F(1, 38) = 9.02$ $p = .0047$ | $F(2.254, 29.30) = 3.79$ $p = .0301$ | $F(7.566, 95.61) = 3.88$ $p = .0007$ | $F(1, 11) = 0.75$ $p = .4057$ | $F(4.970, 54.67) = 1.85$ $p = .1192$ |
| *QW 0-90 min* | | | | | |
| Proportion of time | $F(1, 38) = 116.40$ $p < .0001$ | $F(2.237, 28.34) = 39.22$ $p < .0001$ | – | $t(11) = 2.66$ $p = .0221$ | – |

| | | | | | |
|---|---|---|---|---|---|
| Bout frequency | $F(1, 38) = 41.00$<br>$p < .0001$ | $F(2.210, 27.99) = 14.41$<br>$p < .0001$ | – | $t(11) = 4.21$<br>$p = .0015$ | – |
| Bout duration | $F(1, 38) = 49.79$<br>$p < .0001$ | $F(2.484, 31.46) = 18.47$<br>$p < .0001$ | – | $t(11) = 1.57$<br>$p = .1460$ | – |
| ***NREM sleep*** | | | | | |
| Sleep onset latency | $F(1, 38) < 0.01$<br>$p = .925$ | $F(2.107, 26.69) = 0.06$<br>$p = .945$ | – | $t(11) = 4.57$<br>$p = .0008$ | – |
| Proportion of time | $F(1, 38) = 11.90$<br>$p = .0014$ | $F(2.414, 31.38) = 1.82$<br>$p = .1721$ | $F(8.038, 101.6) = 3.23$<br>$p = .0026$ | $F(1, 11) = 56.21$<br>$p < .0001$ | $F(5.327, 58.60) = 6.80$<br>$p < .0001$ |
| Bout frequency | $F(1, 38) = 0.76$<br>$p = .3877$ | $F(1.921, 24.97) = 0.50$<br>$p = .6073$ | $F(8.652, 109.3) = 1.66$<br>$p = .1111$ | $F(1, 11) = 3.30$<br>$p = .0967$ | $F(4.948, 54.43) = 3.44$<br>$p = .0092$ |
| Bout duration | $F(1, 38) = 0.40$<br>$p = .5305$ | $F(2.286, 29.72) = 0.29$<br>$p = .7822$ | $F(6.848, 86.54) = 1.46$<br>$p = .1955$ | $F(1, 11) = 26.40$<br>$p = .0003$ | $F(4.972, 54.69) = 4.51$<br>$p = .0017$ |
| *NREM sleep 0-90 min* | | | | | |
| Proportion of time | $F(1, 38) = 69.01$<br>$p < .0001$ | $F(2.442, 30.93) = 23.61$<br>$p < .0001$ | – | $t(11) = 8.50$<br>$p < .0001$ | – |
| Bout frequency | $F(1, 38) = 13.74$<br>$p = .0007$ | $F(1.758, 22.26) = 5.33$<br>$p = .0156$ | – | $t(11) = 5.15$<br>$p = .0003$ | – |
| Bout duration | $F(1, 38) = 9.72$<br>$p = .0035$ | $F(2.057, 26.06) = 3.70$<br>$p = .0374$ | – | $t(11) = 6.34$<br>$p < .0001$ | – |
| ***REM sleep*** | | | | | |
| REM sleep onset latency | $F(1, 51) = 105.70$<br>$p < .0001$ | $F(1.898, 32.26) = 35.72$<br>$p < .0001$ | – | $t(11) = 7.55$<br>$p < .0001$ | – |
| Proportion of time | $F(1, 38) = 97.25$<br>$p < .0001$ | $F(2.300, 29.90) = 13.83$<br>$p < .0001$ | $F(8.829, 111.6) = 5.08$<br>$p < .0001$ | $F(1, 11) = 63.97$<br>$p < .0001$ | $F(4.618, 50.80) = 2.93$<br>$p = .0238$ |
| Bout frequency | $F(1, 38) = 36.64$<br>$p < .0001$ | $F(2.234, 29.04) = 9.53$<br>$p = .0005$ | $F(9.211, 116.4) = 3.39$<br>$p = .0009$ | $F(1, 11) = 20.45$<br>$p = .0009$ | $F(4.223, 46.46) = 3.32$<br>$p = .0165$ |
| Bout duration | $F(1, 38) = 29.56$<br>$p < .0001$ | $F(2.053, 26.69) = 9.87$<br>$p = .0006$ | $F(7.762, 98.09) = 5.01$<br>$p < .0001$ | $F(1, 11) = 47.37$<br>$p < .0001$ | $F(5.539, 60.93) = 2.959$<br>$p = .0155$ |
| *REM sleep 30-180 min* | | | | | |
| Proportion of time | $F(1, 38) = 247.90$<br>$p < .0001$ | $F(2.371, 30.03) = 83.28$<br>$p < .0001$ | – | $t(11) = 8.30$<br>$p < .0001$ | – |
| Bout frequency | $F(1, 38) = 169.30$<br>$p < .0001$ | $F(2.335, 29.58) = 57.23$<br>$p < .0001$ | – | $t(11) = 5.46$<br>$p = .0002$ | – |
| Bout duration | $F(1, 38) = 174.50$<br>$p < .0001$ | $F(2.200, 27.86) = 58.70$<br>$p < .0001$ | – | $t(11) = 7.90$<br>$p < .0001$ | – |

**Table 2 – Influence of oxytocin receptor antagonism on i.p. oxytocin-induced effects on sleep-wake outcomes.**

| Sleep-wake outcome | Main effect of dose | Main effect of antagonism | Dose x antagonism interaction |
|---|---|---|---|
| *Active wake (0-30 min)* | | | |
| Proportion of time | $F(1, 3) = 2.12$<br>$p = 0.2414$ | $F(1, 3) = 7.40$<br>$p = 0.0725$ | $F(1, 3) = 1.02$<br>$p = 0.387$ |
| Bout frequency | $F(1, 3) = 9.56$<br>$p = 0.0536$ | $F(1, 3) = 17.70$<br>$p = 0.0245$ | $F(1, 3) = 13.65$<br>$p = 0.0344$ |
| Bout duration | $F(1, 3) = 1.17$<br>$p = 0.3594$ | $F(1, 3) = 0.37$<br>$p = 0.5884$ | $F(1, 3) = 1.12$<br>$p = 0.3677$ |
| *Active wake (30-180 min)* | | | |
| Proportion of time | $F(1, 3) = 5.27$<br>$p = 0.1055$ | $F(1, 3) = 0.01$<br>$p = 0.9207$ | $F(1, 3) = 10.89$<br>$p = 0.0457$ |
| Bout frequency | $F(1, 3) = 3.28$<br>$p = 0.1680$ | $F(1, 3) = 0.45$<br>$p = 0.5504$ | $F(1, 3) = 13.11$<br>$p = 0.0362$ |
| Bout duration | $F(1, 12) = 0.14$<br>$p = 0.7148$ | $F(1, 12) = 2.91$<br>$p = 0.1139$ | $F(1, 12) = 2.20$<br>$p = 0.1641$ |
| *Quiet wake (0-90 min)* | | | |
| Proportion of time | $F(1, 3) = 39.46$<br>$p = 0.0081$ | $F(1, 3) = 3.40$<br>$p = 0.1624$ | $F(1, 3) = 18.30$<br>$p = 0.0235$ |
| Bout frequency | $F(1, 3) = 31.99$<br>$p = 0.0109$ | $F(1, 3) = 0.46$<br>$p = 0.5467$ | $F(1, 3) = 14.68$<br>$p = 0.0313$ |
| Bout duration | $F(1, 3) = 9.79$<br>$p = 0.0521$ | $F(1, 3) = 2.19$<br>$p = 0.2358$ | $F(1, 3) = 6.47$<br>$p = 0.0844$ |
| *NREM sleep (0-60 min)* | | | |
| Sleep onset latency | $F(1, 12) = 1.60$<br>$p = 0.2295$ | $F(1, 12) = 1.88$<br>$p = 0.1951$ | $F(1, 12) = 0.26$<br>$p = 0.6177$ |
| Proportion of time | $F(1, 12) = 44.01$<br>$p = 0.0001$ | $F(1, 12) = 14.56$<br>$p = 0.0025$ | $F(1, 12) = 7.10$<br>$p = 0.0206$ |
| Bout frequency | $F(1, 3) = 6.33$<br>$p = 0.0864$ | $F(1, 3) = 0.67$<br>$p = 0.4726$ | $F(1, 3) = 0.02$<br>$p = 0.903$ |
| Bout duration | $F(1, 3) = 22.47$<br>$p = 0.0178$ | $F(1, 3) = 4.12$<br>$p = 0.1355$ | $F(1, 3) = 1.51$<br>$p = 0.3064$ |
| *REM sleep (30-180 min)* | | | |

| | | | |
|---|---|---|---|
| REM sleep onset latency | $F(1, 3) = 23.95$<br>$p = 0.0163$ | $F(1, 3) = 19.44$<br>$p = 0.0216$ | $F(1, 3) = 72.76$<br>$p = 0.0034$ |
| Proportion of time | $F(1, 3) = 28.91$<br>$p = 0.0126$ | $F(1, 3) = 0.16$<br>$p = 0.7168$ | $F(1, 3) = 9.95$<br>$p = 0.0511$ |
| Bout frequency | $F(1, 3) = 32.93$<br>$p = 0.0105$ | $F(1, 3) = 1.63$<br>$p = 0.291$ | $F(1, 3) = 13.05$<br>$p = 0.0364$ |
| Bout duration | $F(1, 3) = 20.69$<br>$p = 0.0199$ | $F(1, 3) = 0.67$<br>$p = 0.4745$ | $F(1, 3) = 7.17$<br>$p = 0.0751$ |

**Table 3 – Effects of i.n. oxytocin dose range and i.n. caffeine dose on sleep-wake outcomes.**

| Wake outcome | Oxytocin dose range (0, 0.06, 1 mg·kg$^{-1}$) | | | Pairwise comparison | |
|---|---|---|---|---|---|
| | Linear trend | Main effect of dose | Dose x Time interaction | VEH vs 3 mg·kg$^{-1}$ OXT | VEH vs 10 mg·kg$^{-1}$ caffeine |
| ***Active wake*** | | | | | |
| Proportion of time | $F(1, 10) = 0.13$<br>p = .7248 | $F(1.971, 9.857) = 0.15$<br>$p = .8573$ | $F(3.809, 19.04) = 0.80$<br>$p = .5376$ | $t(3) = .083$<br>$p = .9393$ | $t(4) = 6.36$<br>$p = .0031$ |
| Bout frequency | $F(1, 10) < 0.01$<br>$p = .9486$ | $F(1.931, 9.657) = 0.01$<br>$p = .9867$ | $F(3.425, 17.13) = 0.75$<br>$p = .5551$ | $t(3) = 1.60$<br>$p = .2086$ | $t(4) = 5.16$<br>$p = .0067$ |
| Bout duration | $F(1, 10) = 0.84$<br>$p = .3821$ | $F(1.168, 5.842) = 0.71$<br>$p = .4537$ | $F(1.511, 7.553) = 0.97$<br>$p = .3967$ | $t(3) = 1.69$<br>$p = .1892$ | $t(4) = 3.28$<br>$p = .030$ |
| *AW 0-30 min* | | | | | |
| Proportion of time | $F(1, 10) = 1.47$<br>$p = .2534$ | $F(1.630, 8.151) = 0.74$<br>$p = .482$ | – | $t(3) = .44$<br>$p = .6927$ | $t(4) = 5.42$<br>$p = .0056$ |
| Bout frequency | $F(1, 10) = 1.33$<br>$p = .2756$ | $F(1.738, 8.689) = 0.76$<br>$p = .4774$ | – | $t(3) = .78$<br>$p = .4904$ | $t(4) = 2.28$<br>$p = .0845$ |
| Bout duration | $F(1, 10) = 1.65$<br>$p = .2278$ | $F(1.032, 5.162) = 1.02$<br>$p = .3611$ | – | $t(4) = .36$<br>$p = .7415$ | $t(4) = .91$<br>$p = .4146$ |
| *AW 30-180 min* | | | | | |
| Proportion of time | $F(1, 15) < 0.01$<br>$p = .9635$ | $F(1.990, 14.93) = 0.01$<br>$p = .9945$ | – | $t(3) = .01$<br>$p = .993$ | $t(4) = 3.68$<br>$p = .0213$ |
| Bout frequency | $F(1, 15) = 0.16$<br>$p = .6918$ | $F(1.664, 12.48) = 0.15$<br>$p = .8248$ | – | $t(3) = .35$<br>$p = .7479$ | $t(4) = 3.49$<br>$p = .0252$ |
| Bout duration | $F(1, 10) < 0.01$<br>$p = .9934$ | $F(1.588, 7.938) = 0.07$<br>$p = .8966$ | – | $t(3) = 1.48$<br>$p = .235$ | $t(4) = 1.27$<br>$p = .2735$ |
| ***Quiet wake*** | | | | | |
| Proportion of time | $F(1, 10) = 0.22$<br>$p = .6489$ | $F(1.891, 9.454) = 0.12$<br>$p = .8763$ | $F(3.961, 19.81) = 1.35$<br>$p = .2885$ | $t(3) = .45$<br>$p = .6846$ | $t(4) = 1.876$<br>$p = .1339$ |
| Bout frequency | $F(1, 10) = 1.91$<br>$p = .1972$ | $F(1.807, 9.035) = 1.92$<br>$p = .203$ | $F(3.433, 17.17) = 0.69$<br>$p = .5901$ | $t(3) = .09773$<br>$p = .9283$ | $t(4) = 4.519$<br>$p = .0107$ |
| Bout duration | $F(1, 10) = 0.08$<br>$p = .7816$ | $F(1.653, 8.263) = 0.23$<br>$p = .7619$ | $F(3.870, 19.35) = 1.54$<br>$p = .2302$ | $t(3) = .6057$<br>$p = .5875$ | $t(4) = .1051$<br>$p = .9214$ |
| *QW 0-90 min* | | | | | |
| Proportion of time | $F(1, 10) = 9.33$<br>$p = .0121$ | $F(1.645, 8.223) = 4.67$<br>$p = .0491$ | – | $t(3) = .42$<br>$p = .7011$ | $t(4) = 2.12$<br>$p = .1014$ |
| Bout frequency | $F(1, 15) = 5.35$<br>$p = .0354$ | $F(1.703, 12.77) = 2.82$<br>$p = .1026$ | – | $t(3) = .25$<br>$p = .8219$ | $t(4) = 2.78$<br>$p = .0497$ |

| | | | | | |
|---|---|---|---|---|---|
| Bout duration | $F(1, 10) = 2.00$<br>$p = .1878$ | $F(1.766, 8.832) = 1.64$<br>$p = .2465$ | – | $t(3) = 1.48$<br>$p = .235$ | $t(4) = 1.27$<br>$p = .2735$ |
| ***NREM sleep*** | | | | | |
| Sleep onset latency | $F(1, 10) = 0.03$<br>$p = .8758$ | $F(1.793, 8.966) = 0.04$<br>$p = .9452$ | – | $t(3) = 0.99$<br>$p = .396$ | $t(4) = 1.92$<br>$p = .1267$ |
| Proportion of time | $F(1, 10) = 1.48$<br>$p = .2515$ | $F(1.517, 7.583) = 0.98$<br>$p = .3923$ | $F(3.593, 17.97) = 0.52$<br>$p = .7043$ | $t(3) = .43$<br>$p = .6955$ | $t(4) = 7.03$<br>$p = .0022$ |
| Bout frequency | $F(1, 10) = 0.15$<br>$p = .7072$ | $F(1.184, 5.920) = 0.10$<br>$p = .8038$ | $F(4.092, 20.46) = 0.57$<br>$p = .6905$ | $t(3) = .63$<br>$p = .5728$ | $t(4) = 2.54$<br>$p = .064$ |
| Bout duration | $F(1, 10) = 0.46$<br>$p = .5145$ | $F(1.434, 7.168) = 0.24$<br>$p = .7253$ | $F(3.462, 17.31) = 0.68$<br>$p = .5966$ | $t(3) = .21$<br>$p = .8473$ | $t(4) = 6.46$<br>$p = .003$ |
| *NREM sleep 0-90 min* | | | | | |
| Proportion of time | $F(1, 10) = 1.34$<br>$p = .2735$ | $F(1.857, 9.285) = 0.76$<br>$p = .4839$ | – | $t(3) = .08$<br>p = .9378 | $t(4) = 4.36$<br>p = .012 |
| Bout frequency | $F(1, 15) = 0.31$<br>$p = .5875$ | $F(1.979, 14.84) = 0.17$<br>$p = .8446$ | – | $t(3) = .65$<br>$p = .5595$ | $t(4) = 1.74$<br>$p = .1571$ |
| Bout duration | $F(1, 10) = 0.28$<br>$p = .6059$ | $F(1.810, 9.051) = 0.21$<br>$p = .7918$ | – | $t(3) = .17$<br>$p = .8740$ | $t(4) = 7.13$<br>$p = .0020$ |
| ***REM sleep*** | | | | | |
| REM sleep onset latency | $F(1, 15) = 0.03$<br>$p = .8747$ | $F(1.445, 10.84) = 1.60$<br>$p = .2416$ | – | $t(3) = .75$<br>$p = .5087$ | $t(4) = 2.15$<br>$p = .0975$ |
| Proportion of time | $F(1, 10) = 1.79$<br>$p = .2111$ | $F(1.289, 6.446) = 1.05$<br>$p = .3657$ | $F(4.114, 20.57) = 1.36$<br>$p = .2815$ | $t(3) = .6323$<br>$p = .5721$ | $t(4) = 5.57$<br>$p = .0051$ |
| Bout frequency | $F(1, 10) = 0.48$<br>$p = .5054$ | $F(1.292, 6.460) = 0.190$<br>$p = .7366$ | $F(4.094, 20.47) = 0.43$<br>$p = .7925$ | $t(3) = .89$<br>$p = .4409$ | $t(4) = 2.49$<br>$p = .0674$ |
| Bout duration | $F(1, 10) = 3.14$<br>$p = .1067$ | $F(1.265, 6.325) = 2.21$<br>$p = .1876$ | $F(3.901, 19.51) = 1.30$<br>$p = .3060$ | $t(3) = 0.75$<br>$p = .5054$ | $t(4) = 4.28$<br>$p = .0128$ |
| *REM sleep 30-180 min* | | | | | |
| Proportion of time | $F(1, 15) = 0.64$<br>$p = .4375$ | $F(1.767, 13.25) = 0.58$<br>$p = .5534$ | – | $t(3) = .23$<br>$p = .834$ | $t(4) = 3.89$<br>$p = .0176$ |
| Bout frequency | $F(1, 10) = 1.54$<br>$p = .2432$ | $F(1.726, 8.631) = 0.77$<br>$p = .4745$ | – | $t(3) = .93$<br>$p = .4208$ | $t(4) = 2.91$<br>$p = .0436$ |
| Bout duration | $F(1, 15) = 0.36$<br>$p = .5555$ | $F(1.671, 12.53) = 0.33$<br>$p = .6863$ | – | $t(3) = .63$<br>$p = .5723$ | $t(4) = 3.25$<br>$p = .0315$ |
