## Supplemental material for "Quiet wakefulness: The influence of intraperitoneal and intranasal oxytocin on sleep-wake behaviour and neurophysiology in rats"

#### Methods

##### *Animals and housing*

Eight-week-old male and female Wistar rats (ARC, WA, Australia), weighing on average 383 g (M) and 241 g (F) upon arrival. This species and strain were chosen as Wistar rats are the most common species/strain used in previous preclinical research on oxytocin-based interventions and sleep[1]. Rats were housed in pairs in filter-top cages (size: 58 x 38 x 20 cm; Able Scientific) containing corn cob bedding material (Bed-o' Cobs, The Andersons) and environmental enrichment including shredded paper nesting material (Crink-l'Nest, The Andersons) and two wooden gnawing logs. All experiments were conducted within a specific pathogen free (SPF) facility and rats were housed in a temperature- and humidity-controlled room (22 ± 0.5°C; 50-60%) under a reverse light cycle (12L:12D; lights on at 1400, ZT0). At transitions between light phases, illumination was slowly transitioned between 0 and 500 lux over a 16-min period. Rats had *ad libitum* access to standard rodent chow and water at all times during the experiment. Prior to commencing experimental procedures, all rats were allowed at least 3 days to acclimatise to the facility.

General welfare monitoring for rats was conducted at least twice per week including assessments of body weight, gait and posture, hydration, and grooming. Acute monitoring during surgery involved regular assessment of anaesthetic levels and oxygen flow rate, withdrawal reflex, respiration rate, and body temperature. Post-operative monitoring was conducted on a daily basis with a particular emphasis on assessing pain Rat Grimace Scale; [2]. Pre-defined specific euthanasia criteria (humane endpoints) included: gasping respiration, low response to stimuli, emaciation, loss of >15% body weight, and reopening of surgical incisions and irreparable damage to probe wiring. Euthanasia was conducted via i.p. injection of sodium pentobarbital (~100 $mg \cdot kg^{-1}$). All experiments were conducted in line with the *Australian code for the care and use of animals for scientific purposes (8th edition, 2013)* and were approved by the Animal Ethics Committee at The University of Sydney (AEC number: 2019/1615).

##### *Drug preparation*

Oxytocin (China Peptide, China) and caffeine (anhydrous; AK Scientific Inc., USA) were dissolved in a vehicle of physiological saline (0.9% w/v). The non-peptidergic oxytocin receptor (OXTR) antagonist L-368,899 hydrochloride (Santa Cruz Biotechnology, Texas) was dissolved in a vehicle of dimethylsulfoxide (DMSO; 5% v/v) Tween 80 (5% v/v) and saline (90% v/v).

##### *Intraperitoneal administration*

Oxytocin was administered at light onset (ZT0) via intraperitoneal (I.P.) injection at a range of doses (0.1, 0.3, and 1 $mg \cdot kg^{-1}$ b.w.) at an injection volume of 1 $mL \cdot kg^{-1}$ body weight. These doses were selected to cover doses of oxytocin that do (1 mg/kg) and do not (0.3 and 0.1 mg/kg) have significant effects on gross locomotor activity in rats [3]. Caffeine (I.P.) was administered at ZT0 at a dose of 10 $mg \cdot kg^{-1}$ as a positive control based on previous research demonstrating potent wake-promoting effects at this dose in rats [4,5]. Pre-treatment using the antagonist L-368,899 (5 mg·kg$^{-1}$) was conducted I.P. 15-min prior to light onset (ZT23.75). The OXTR antagonist L-368,899 was chosen as systemic administration of L-368,899 is centrally penetrant and preferentially binds to OXTRs [6]. The 5 mg·kg$^{-1}$ dose of L-368,899 was chosen based on pre-treatment of this dose blocking the stress-attenuating effects of 1 mg·kg$^{-1}$ oxytocin in rats[7].

***Intranasal administration***

Oxytocin (0.06, 1, and 3 mg·kg$^{-1}$) and caffeine (10 mg·kg$^{-1}$) were administered intranasally (i.n.) according to methods adapted from Lukas and Neumann [8]. In brief, unanaesthetised rats were restrained by an experienced experimenter and the solution was applied bilaterally to the rhinarium (10 uL to each side) using a 12.5 µL automated pipette by another experimenter, allowing approximately 1 minute for the solution to diffuse into the nasal epithelium. The 0.06 mg·kg$^{-1}$ dose represents a ten-fold higher dose than the highest i.n. dose administered in human clinical research 80 IU; [9]; the human equivalent dose of 0.06 mg·kg$^{-1}$ is ~774 IU for an 80 kg human based on FDA guidelines of rat to human dose conversion; [10]. This dose was selected to investigate whether doses closer to—albeit still substantially higher than—those administered in clinical research would impact sleep-wake behaviour. The 1 and 3 mg·kg$^{-1}$ doses were chosen to facilitate comparisons between i.p. and i.n. routes of administration; the higher dose was based on a previous study demonstrating that i.p. oxytocin elicited approximately two-fold greater increases in plasma and central oxytocin levels compared to the same dose administered i.n.[11].

***Radiotelemetry probe implantation surgery***

Rats were surgically implanted with wireless radiotelemetry probes capable of polysomnographic (PSG)—electrocorticographic (ECoG) and electromyographic (EMG)—recording (HD-X02, Data Sciences International Inc.). Surgery was conducted using aseptic technique and isoflurane gas anaesthesia (3% and 1.5-2% in 1.2 L·min$^{-1}$ for induction and maintenance, respectively). Rats were administered buprenorphine as a pre-operative analgesic (0.05 mg·kg$^{-1}$; s.c.) and bupivacaine locally at incision locations (0.25% w/v), on the flank and above skull. This was followed by mounting the rat onto the stereotaxic apparatus (Kopf, CA, USA), inserting a rectal thermal probe to monitor temperature, and placing a heat pad underneath to ensure a constant body temperature. The fur covering the flank and skull was removed and the incision sites were cleaned and sterilised using both povidone-iodine (Riodine, WA, Australia) and ethanol (80% v/v) antiseptics. Briefly, after induction and administration of pre-operative analgesics, rats were placed into the stereotaxic apparatus (Kopf, CA, USA). An incision was made above the right flank, a telemetry probe was implanted subcutaneously under this incision, and the probe wires were fed under the skin towards the head. Another incision was made above the skull and two holes (1-mm diameter) were drilled through the skull in fronto-parietal locations (measured from bregma: (1) anterior/posterior: +2 mm, lateral: +1.5 mm; (2) anterior/posterior: –7 mm, lateral: –1.5 mm). Screws were inserted into these holes until they contacted the dura, and then ECoG wires were wrapped around the screws and secured using Vetbond (3M, USA) and Kwik-Sil (World Precision Instruments, USA) to isolate electrical conductance to the dura. EMG wires were partially stripped of insulation to reveal bare wire, threaded through trapezius muscle behind the skull, and then insulation was replaced and secured to ensure electrical conductance was isolated to the muscle. Surgical incisions were sutured back together, and rats received post-operative analgesia, non-steroidal anti-inflammatory meloxicam (2 mg·kg$^{-1}$; s.c.), after the surgery and for the two subsequent days as a post-operative analgesia. Rats were allowed at least 14 days recovery prior to experimentation.

***Habituation***

Rats were handled across the seven days prior to surgery and on the days following the recovery period. Habituation i.p. saline injections were administered on three of these days. Habituation to the telemetry recording cages and procedure was also conducted with three 6-hour baseline PSG recordings performed prior to testing to ensure proper probe functioning and reception by telemetry receiver plates. Two habituation sessions involving restraint and nasal administration of saline were conducted prior to i.n. experimentation to familiarise rats to the i.n. administration procedure.

***Polysomnographic recordings***

On recording days, probes were turned on by holding a magnet close to the rat's flank (above the probe) and then rats were placed individually into telemetry cages (43 x 26 x 12 cm, Techniplast, Italy) located above telemetry receiver plates (RPC-1, Data Sciences International Inc.). For all experiments except Experiment 2, rats were placed into recording cages at least 1 hour prior to light onset and recordings were started by 1300 (ZT23). Due to pre-treatment with antagonists for Experiment 2, rats were placed into recording cages at least 1.25 hour prior to light onset and recordings were started by 1245 (ZT22.75). At light onset (ZT0), rats were removed from the telemetry cage, administered an i.p. injection or i.n. application, placed back in the cage, left to behave freely for 6 hours (until ZT6), and then returned to their home cages (Figure 1A). For female rats, oestrus phase determination was conducted according to methods adapted from Marcondes, Bianchi and Tanno [12], immediately following the termination of PSG recording sessions (ZT6-6.5).

***Experimental design***

All experiments were conducted across consecutive recording sessions using repeated-measures, counterbalanced designs to minimise inter-subject variability and increase statistical power. In all designs, dose sequences were generated using William's Latin Square designs that control for first-order carryover effects [13], and rats were randomised to dose sequences using a random number generator, with the caveat that pair-housed rats could not be assigned to the same dose sequence. There was at least 90-h washout before the next recording session following oxytocin or L-368,899 administration, and 7 days following caffeine, which demonstrates more marked and prolonged effects on sleep [5,14]. Hence, the caffeine condition was not incorporated into oxytocin dose-response William's Latin square designs in Experiment 1 to preserve a consistent washout period between recording sessions.

*Oxytocin i.p. dose-response (Experiments 1A and 1B)*

This experiment characterised the effects of i.p. oxytocin (VEH, 0.1, 0.3, 1 mg·kg$^{-1}$; Experiment 1A) and caffeine (VEH and 10 mg·kg$^{-1}$; Experiment 1B) on sleep-wake outcomes. Male and female rats ($N$ = 16; $n$ = 8 per sex) were run as separate cohorts to avoid potential interference of opposite-sex pheromones in sleep behaviour and physiology during PSG recording sessions [15].

*Oxytocin i.p. and OXTR antagonism (Experiment 2)*
This experiment explored whether i.p. oxytocin-induced effects on sleep-wake outcomes are mediated by the OXTR. Since no significant sex by dose interactions were apparent during Experiment 1 (See Supplementary Material Table S1), only female rats ($n$ = 4) were used for Experiment 2. The following conditions examined whether an OXTR antagonist inhibited oxytocin effects on sleep-wake outcomes: (i) VEH + VEH; (ii) VEH + oxytocin (1 mg·kg$^{-1}$); (iii) L-368,899 (5 mg·kg$^{-1}$) + VEH; (iv) L-368,899 (5 mg·kg$^{-1}$) + oxytocin (1 mg·kg$^{-1}$).

*Oxytocin i.n. dose-response (Experiments 3A and 3B)*
Experiment 3A characterised the dose-dependent effects of i.n. oxytocin (0, 0.06, 1 mg·kg$^{-1}$ in 20 μL), and Experiment 3B a higher dose of i.n. oxytocin (3 mg·kg$^{-1}$ in 20 μL) and caffeine (10 mg·kg$^{-1}$ in 20 μL), on sleep-wake outcomes. The goal was to examine whether i.n. oxytocin would recapitulate the effects of i.p. oxytocin observed in Experiment 1. As with Experiment 2, female rats were used: $n$ = 6 for Experiment 3A and $n$ = 5 for Experiment 3B.

***Oestrus phase determination***
Briefly, while the rat was restrained, the tip of a plastic Pasteur pipette containing ~0.1-0.2 mL of saline (0.9 % w/v) was carefully inserted into the vagina (≤ 1 cm insertion depth), the saline was expelled to gently flush the vagina, and then vaginal fluid was collected into the pipette. Vaginal fluid (2-3 drops) from each rat was placed onto individual labelled glass slides and then unstained samples were examined under a light microscope using 10 and 40 x objective lenses. Additionally, photographs (2-3 images) of each sample were taken using a microscope digital camera (M500 BASE, Levenhuk) and software (ToupView, Levenhuk). Oestrus phase was determined based on the relative proportion of cell types (epithelial, cornified and leukocytes) present in the sample: high prevalence of nucleated epithelial cells in proestrus phase; high prevalence of enucleated cornified epithelial cells in oestrus phase; roughly equal prevalence of epithelial (both nucleated and cornified) and leukocytes in metoestrus phase, and high prevalence of leukocytes in dioestrus phase [16]. After oestrus phase determination, rats were returned to their home cages and glass slides were cleaned thoroughly and allowed to dry.

***Data acquisition, processing, and analysis***
Polysomnographic recording data (ECoG, EMG, body temperature and activity) from freely moving rats were transmitted from telemetry probes to receiver plates connected to a matrix 2.0 (MX-2) and relayed to a computer located outside of the testing room. Data were acquired using Ponemah software (Version 6.41, Data Sciences International). NeuroScore (Version 3.2.1, Data Sciences International) was used to score sleep-wake states for each 10-s

epoch of time: active wake (AW), quiet wake (QW), non-REM sleep (NREMS) and REM sleep (REMS). Pre-REM sleep, the transition state between NREM and REM sleep characterised by high amplitude EEG activity within the theta and alpha power bands (Benington et al., 1994), was scored as NREM sleep. ECoG and EMG signals were filtered prior to scoring: a band-pass filter (0.1-80 Hz) and high-pass filter (> 0.1 Hz) were applied to the ECoG and EMG signal, respectively. Sleep scoring was conducted manually by a blinded experimenter via visual examination of various raw and derived signals: power spectral density 'periodogram' of ECoG signal (Fast Fourier Transform; 0-25 Hz), theta:delta ratio, ECoG trace, EMG trace, activity counts, and sleep-wake state of previous epoch. Briefly, scoring criteria for each sleep-wake state were as follows (see Figure 1B for representative traces):

- *Active wake*: (ECoG) low amplitude, low synchrony, high frequency; (EMG) high amplitude, irregular tone; (Activity) locomotion and movement present
- *Quiet wake*: (ECoG) low amplitude, low synchrony, moderate-high frequency relative to AW; (EMG) moderate amplitude relative to AW, regular tone; (Activity) no locomotion and minimal movement
- *NREM sleep*: (ECoG) moderate-high amplitude, high synchrony, very low-low frequency; (Periodogram) high power density within the delta frequency band (0-4 Hz); (EMG) low amplitude, regular tone; (Activity) no locomotion or movement
- *REM sleep*: (ECoG) low amplitude, high frequency, 'saw-tooth' profile; (Periodogram) high power density within the theta frequency band (4-8 Hz) and high theta:delta ratio; (EMG) low amplitude, regular tone; (Activity) no locomotion or movement
- *Artefact*: epochs were scored as artefacts and excluded from analysis if no signals (ECoG, EMG, activity etc.) were available during the epoch to indicate sleep-wake state.

After sleep scoring was completed, the following sleep architectural outcomes of interest were extracted: sleep onset latency (min), REM sleep onset latency (min), proportion of total time spent in each sleep state (%), bout frequency of each sleep state, mean duration of bout of each sleep state (min), and mean body temperature (ºC). These 7 hours of PSG data (1-h prior to oxytocin administration during last hour of dark phase and 6 hours post-administration during light phase) were then compiled into 30-min bins for statistical analysis. For complete details on subject attrition and data exclusion, see Supplemental materials.

ECoG power spectral density outcomes during NREM sleep, QW, and REM sleep were extracted for each 1-Hz frequency band from 0-25 Hz using Discrete Fourier Transform (DFT) and Hamming window function. Automated artefact detection was conducted using an absolute threshold exclusion criteria: if ECoG signals were equal to or exceeded an amplitude of 0.5 mV during a 10-s epoch, these epochs were excluded from analysis. Subsequently, data were manual screened for artefacts where ECoG signals were absent (i.e., 0 mV) and these were also excluded. As large inter-individual differences existed in ECoG spectral outcomes, all subjects' data were normalised by expressing values as a proportion of an outcome during the corresponding VEH condition of each respective experiment. To elaborate, spectral density ($V^2$) was averaged over 0-1 h and 1-6 h bins of the recording during the light phase for each sleep-wake state (NREM, REM and QW), for each 1-Hz frequency bin from 0-25 Hz, for each dose; expressed as a proportion of this average during the VEH recording (%); and then the percentage change (%Δ) was calculated.

As sleep outcomes appear to be predominantly influenced by the oestrus cycle during proestrus and oestrus phases [17,18], oestrus stage was delineated into two categories: either proestrus/oestrus or metoestrus/dioestrus.

***Statistical analysis***

Linear mixed effects models (LMMs) were constructed for all sleep architectural outcomes with *Dose, Time,* and *Dose* x *Time* as the within-subjects fixed effects factors. LMMs were chosen to avoid having to exclude subjects with missing data. A compound symmetry covariance matrix structure was used and fitted using restricted maximum likelihood estimation. Greenhouse-Geisser corrections were applied to adjust for violations of sphericity. If random effects were zero, they were removed from the model and a simpler model was fitted.

For sleep architectural outcomes, the first hour of data (dark phase; pre-administration) were analysed separately from the following 6 hours (light phase; post-administration) to test for baseline differences in outcomes. As part of the LMM analysis, trend analysis was conducted on *Dose* to determine if a linear dose-dependent effect was present. Based on the pharmacokinetics of acute peripherally administered oxytocin ($t_{1/2}$ = < 5 min; Troncy, Morin, del Castillo, Authier, Ybarra, Otis, Gauvin and Gutkowska [19]) and *post hoc* visual inspection of the sleep architectural outcomes, average values were calculated for periods of the peak effect for each outcome—AW: 0-30 min and 30-180 min, QW: 0-90 min, NREM sleep: 0-90 min, REM sleep: 30-180 min, and body temperature: 0-120 min. LMM and trend analyses were conducted on these composite outcomes with *Dose* as a fixed effect.

For Experiment 1A, data were analysed as pre-administration (2 x 4) and post-administration (12 x 4) LMMs including *Time* and *Dose* as fixed effects. For Experiment 1B, data were analysed as pre-administration (2 x 2) and post-administration (12 x 2) LMMs including *Time* and *Dose* as fixed effects. For Experiment 2, composite scores were analysed as a 2 x 2 LMM including *Oxytocin* and *Antagonist* as fixed effects. For Experiments 3A and 3B, data were analysed as pre-administration (2 x 3) and post-administration (12 x 3) LMMs including *Time* and *Dose* as fixed effects. However, while trend analysis was conducted for Experiment 3A, planned contrasts were conducted for Experiment 3B comparing 3 $mg \cdot kg^{-1}$ oxytocin and 10 $mg \cdot kg^{-1}$ Caffeine to VEH using two-sided paired t-tests.

For ECoG power spectral density outcomes, normalised average values (%) for each 1-Hz frequency band were transformed using a logarithmic ($\log_{10}$) transformation, and then compared to the baseline (VEH) value using a two-tailed one-sample t-test. For oestrus phase data, differences in the proportion of rats in proestrus/oestrus and metoestrus/dioestrus phases during each experiment between each dose and VEH were analysed using two-sided Fisher's

Exact tests. Fisher's least significant difference (LSD) test was used to control Type I error rate. Statistical analyses were conducted using GraphPad Prism (Version 9.3.1). The level of significance for all tests was $p < .05$.

#### *Attrition and exclusion of data*

Details of attrition and data exclusion are reported here for each experiment. In Experiment 1A, data were excluded from $n = 1$ female subject during the 0.1 $mg \cdot kg^{-1}$ oxytocin dose recording due to complete signal dropout from the telemetry probe to the receiver. Sleep scoring data from all dose recordings were excluded for $n = 1$ male subject due to signal interference between ECoG and EMG signals that meant scoring could not be accurately conducted, however temperature measurement was unaffected. Additionally, $n = 1$ male subjects was euthanised due to post-surgical complications. In Experiment 1B, $n = 2$ female subjects were euthanised due to complications involving rats creating lesions to access and damage insulation of probe wires. In Experiment 3B, sleep scoring data for $n = 1$ female subject were excluded during the 3.0 $mg \cdot kg^{-1}$ oxytocin dose recording due to signal dropout.

#### *Sex difference statistical analyses*

Preliminary statistical analyses were conducted to determine if sex differences existed in the response to difference doses of oxytocin across primary sleep-wake outcomes in Experiment 1a (i.e., represented by significant sex x dose interaction effects). Analyses were conducted using 2 x 4 (sex x dose) mixed-model analysis on aggregate measures of proportion of time spent in sleep-wake states and onset latencies.

### Results

#### *Experiment 1*

*Sex difference data*

Preliminary analyses of data from Experiment 1a (oxytocin i.p. dose-response experiment), revealed some significant main effects of sex: on average, males experienced a greater proportion of time in QW during 0-90 min and in REM sleep during 30-180 min than females, and females experienced a greater proportion of time in NREM sleep during 0-90 min than males (see highlighted values in Table S1). However, no significant *Dose* x *Sex* interaction effects were observed across any primary sleep-wake outcomes. Hence, it was decided that data from both sexes would be combined for analyses in Experiment 1, and that Experiments 2 and 3 would be conducted solely in female rats.

**Table S1 – Preliminary analyses of sex effects in oxytocin i.p. dose-response experiment.**

| Outcome | Sex (main effect) | p-value | Sex x Dose interaction | p-value |
|---|---|---|---|---|
| ***Active wake*** | | | | |
| %AW 0-30 min | $F(1, 12) = 1.72$ | $p = .2146$ | $F(3, 35) = 1.09$ | $p = .3682$ |
| %AW 30-180 min | $F(1, 12) = 1.97$ | $p = .1856$ | $F(3, 35) = 0.54$ | $p = .6558$ |
| ***Quiet wake*** | | | | |
| %QW 0-90 min | $F(1, 12) = 5.28$ | $p = .0403$ | $F(3, 35) = 0.39$ | $p = .7577$ |
| ***NREM sleep*** | | | | |

| | | | | |
|---|---|---|---|---|
| %NREM 0-90 min | $F(1, 12) = 6.18$ | $p = .0286$ | $F(3, 35) = 0.36$ | $p = .7791$ |
| SOL | $F(1, 12) = 1.20$ | $p = .2950$ | $F(3, 35) = 0.33$ | $p = .8040$ |
| ***% REM sleep*** | | | | |
| %REM 30-180 min | $F(1, 12) = 5.36$ | $p = .0392$ | $F(3, 35) = 0.08$ | $p = .9682$ |
| ROL | $F(1, 47) = 2.11$ | $p = .1533$ | $F(3, 47) = 0.73$ | $p = .5390$ |

*Power spectral density data*

*Quiet wake:* During the 1st hour post-administration, relative to vehicle, 1 mg·kg$^{-1}$ oxytocin reduced ECoG PSD in the theta, alpha and sigma frequency bands, while caffeine reduced ECoG PSD in the delta, alpha, sigma, and beta frequency bands. In contrast, during the last 5 hours of recording, oxytocin increased ECoG PSD in the delta (0.1 and 1 mg·kg$^{-1}$), theta (all doses), and alpha (0.1 mg·kg$^{-1}$) frequency bands, whereas caffeine reduced ECoG PSD within the alpha, sigma, and beta frequency bands, relative to vehicle.

*NREM sleep*: During the 1st hour post-administration, relative to vehicle, 1 mg·kg$^{-1}$ oxytocin reduced ECoG PSD in the theta, alpha, sigma, and beta frequency bands, while caffeine reduced ECoG PSD across all frequency bands. In contrast, during the last 5 hours of recording, oxytocin increased ECoG PSD in the delta (all doses), theta (all doses), alpha (all doses), and beta (0.1 mg·kg$^{-1}$) frequency bands, whereas caffeine had no effect on ECoG PSD.

*REM sleep*: Over the 6-hour recording post-administration, relative to vehicle, oxytocin increased ECoG PSD in the delta (0.1 and 1 mg·kg$^{-1}$), alpha (all doses), sigma (0.1 and 1 mg·kg$^{-1}$), and beta (all doses) frequency bands. In contrast, caffeine reduced ECoG PSD in the sigma and beta frequency bands.

**Table S2 – Effects of i.p. oxytocin dose range and i.p. caffeine dose on power spectral density outcomes during quiet wake, NREM sleep and REM sleep.**

| Relative to VEH | Oxytocin (mg·kg$^{-1}$) | | | Caffeine (mg·kg$^{-1}$) |
|---|---|---|---|---|
| | ***0.1*** | ***0.3*** | ***1.0*** | ***10*** |
| ***QW (0-1 h)*** | | | | |
| *Delta (0.1-4 Hz)* | $t(12) = 1.57$ $p = .1423$ | $t(13) = 0.14$ $p = .8917$ | $t(13) = 0.05$ $p = .9647$ | $t(11) = 2.37$ $p = .0374$ |
| *Theta (4-8 Hz)* | $t(12) = 1.08$ $p = .3025$ | $t(13) = 1.18$ $p = .2577$ | $t(13) = 2.79$ $p = .0152$ | $t(11) = 1.39$ $p = .1917$ |
| *Alpha (8-12 Hz)* | $t(12) = 0.39$ $p = .7023$ | $t(13) = 2.11$ $p = .0545$ | $t(13) = 2.64$ $p = .0206$ | $t(11) = 3.54$ $p = .0046$ |
| *Sigma (12-16 Hz)* | $t(12) = 0.39$ $p = .7011$ | $t(13) = 1.55$ $p = .1449$ | $t(13) = 2.64$ $p = .0206$ | $t(11) = 5.53$ $p = .0002$ |
| *Beta (16-25 Hz)* | $t(12) = 1.18$ $p = .2611$ | $t(13) = 0.22$ $p = .8321$ | $t(13) = 1.33$ $p = .2052$ | $t(11) = 4.39$ $p = .0011$ |
| ***QW (1-6 h)*** | | | | |
| *Delta (0.1-4 Hz)* | $t(12) = 2.46$ $p = .0302$ | $t(13) = 1.84$ $p = .0885$ | $t(13) = 3.37$ $p = .0050$ | $t(11) = 2.17$ $p = .0533$ |
| *Theta (4-8 Hz)* | $t(12) = 2.53$ $p = .0266$ | $t(13) = 3.21$ $p = .0069$ | $t(13) = 3.56$ $p = .0035$ | $t(11) = 1.96$ $p = .0760$ |
| *Alpha (8-12 Hz)* | $t(12) = 2.19$ $p = .0490$ | $t(13) = 1.33$ $p = .2056$ | $t(13) = 1.85$ $p = .0867$ | $t(11) = 4.09$ $p = .0018$ |
| *Sigma (12-16 Hz)* | $t(12) = 0.76$ $p = .4626$ | $t(13) = 0.29$ $p = .7794$ | $t(13) = 0.40$ $p = .6930$ | $t(11) = 6.17$ $p < .0001$ |
| *Beta (16-25 Hz)* | $t(12) = 1.88$ | $t(13) = 0.96$ | $t(13) = 1.79$ | $t(11) = 5.76$ |

| | | | | |
|---|---|---|---|---|
| | $p = .0849$ | $p = .3548$ | $p = .0971$ | $p = .0001$ |
| ***NREM (0-1 h)*** | | | | |
| *Delta (0.1-4 Hz)* | $t(12) = 2.07$<br>$p = .0611$ | $t(13) = 0.07$<br>$p = .9484$ | $t(13) = 1.87$<br>$p = .0846$ | $t(7) = 5.12$<br>$p = .0014$ |
| *Theta (4-8 Hz)* | $t(12) = 2.08$<br>$p = .0598$ | $t(13) = 0.14$<br>$p = .8913$ | $t(13) = 2.37$<br>$p = .0340$ | $t(7) = 4.34$<br>$p = .0034$ |
| *Alpha (8-12 Hz)* | $t(12) = 1.64$<br>$p = .1260$ | $t(13) = 1.12$<br>$p = .2818$ | $t(13) = 3.48$<br>$p = .0041$ | $t(7) = 3.45$<br>$p = .0108$ |
| *Sigma (12-16 Hz)* | $t(12) = 0.34$<br>$p = .7429$ | $t(13) = 1.35$<br>$p = .2010$ | $t(13) = 4.06$<br>$p = .0013$ | $t(7) = 4.03$<br>$p = .0050$ |
| *Beta (16-25 Hz)* | $t(12) = 1.71$<br>$p = .1129$ | $t(13) = 0.53$<br>$p = .6028$ | $t(13) = 2.66$<br>$p = .0195$ | $t(7) = 4.34$<br>$p = .0034$ |
| ***NREM (1-6 h)*** | | | | |
| *Delta (0.1-4 Hz)* | $t(12) = 3.58$<br>$p = .0038$ | $t(13) = 3.50$<br>$p = .0039$ | $t(13) = 4.98$<br>$p = .0003$ | $t(11) = 2.05$<br>$p = .0652$ |
| *Theta (4-8 Hz)* | $t(12) = 4.27$<br>$p = .0011$ | $t(13) = 4.40$<br>$p = .0007$ | $t(13) = 4.69$<br>$p = .0004$ | $t(11) = 1.25$<br>$p = .2392$ |
| *Alpha (8-12 Hz)* | $t(12) = 3.17$<br>$p = .0081$ | $t(13) = 2.37$<br>$p = .0343$ | $t(13) = 2.67$<br>$p = .0193$ | $t(11) = 1.08$<br>$p = .3051$ |
| *Sigma (12-16 Hz)* | $t(12) = 0.78$<br>$p = .4505$ | $t(13) = 0.61$<br>$p = .5500$ | $t(13) = 0.23$<br>$p = .8234$ | $t(11) = 0.75$<br>$p = .4700$ |
| *Beta (16-25 Hz)* | $t(12) = 2.65$<br>$p = .0212$ | $t(13) = 1.86$<br>$p = .0858$ | $t(13) = 1.69$<br>$p = .1143$ | $t(11) = 0.87$<br>$p = .4036$ |
| ***REM (0-6 h)*** | | | | |
| *Delta (0.1-4 Hz)* | $t(12) = 2.72$<br>$p = .0188$ | $t(13) = 2.13$<br>$p = .0526$ | $t(13) = 3.02$<br>$p = .0099$ | $t(11) = 0.96$<br>$p = .3578$ |
| *Theta (4-8 Hz)* | $t(12) = 0.35$<br>$p = .7331$ | $t(13) = 2.07$<br>$p = .0586$ | $t(13) = 1.81$<br>$p = .0937$ | $t(11) = 0.27$<br>$p = .7898$ |
| *Alpha (8-12 Hz)* | $t(12) = 2.86$<br>$p = .0144$ | $t(13) = 2.55$<br>$p = .0241$ | $t(13) = 3.64$<br>$p = .0030$ | $t(11) = 1.61$<br>$p = .1363$ |
| *Sigma (12-16 Hz)* | $t(12) = 2.09$<br>$p = .0587$ | $t(13) = 2.35$<br>$p = .0350$ | $t(13) = 4.20$<br>$p = .0010$ | $t(11) = 4.78$<br>$p = .0006$ |
| *Beta (16-25 Hz)* | $t(12) = 2.49$<br>$p = .0284$ | $t(13) = 2.95$<br>$p = .0113$ | $t(13) = 3.59$<br>$p = .0033$ | $t(11) = 5.63$<br>$p = .0002$ |

*Oestrus data*

No significant differences were found in the proportion of rats in proestrus/oestrus and metoestrus/dioestrus stages between oxytocin dose conditions and VEH in Experiment 1a (Table S3). Additionally, no significant difference in the proportion of rats in proestrus/oestrus and metoestrus/dioestrus stages between caffeine and VEH was observed in Experiment 1b (Table S4).

**Table S3 – Proportion of rats in oestrus phases for each dose condition in Experiment 1a.**

| Oxytocin dose (mg·kg$^{-1}$) | Proestrus/Oestrus (%) | Metoestrus/Dioestrus (%) | Fisher's exact test (2-sided) |
|---|---|---|---|
| **VEH** | 0 | 100 | |
| **0.1** | 0 | 100 | $p > .9999$ |
| **0.3** | 12.5 | 87.5 | $p > .9999$ |
| **1.0** | 12.5 | 87.5 | $p > .9999$ |

**Table S4 – Proportion of rats in oestrus phases for each dose condition in Experiment 1b.**

| Caffeine dose (mg·kg$^{-1}$) | Proestrus/Oestrus (%) | Metoestrus/Dioestrus (%) | Fisher's exact test (2-sided) |
|---|---|---|---|
| **VEH** | 16.7 | 83.3 | |
| **10** | 16.7 | 83.3 | $p > .9999$ |

*Body temperature data*

For the oxytocin i.p. dose response experiment (Experiment 1a), no significant main effect of dose [$F(1.490, 20.86) = 0.80$, $p = .4279$] or interaction effect of dose x time [$F(2.54, 33.97) = 0.28$, $p = .8064$] were observed during baseline recording (pre-dose) for the last hour of the dark phase. Across the 6 hours following administration of i.p. oxytocin, a significant main effect of dose [$F(2.378, 33.29) = 12.74$, $p < .0001$] and a significant interaction effect of dose x time [$F(6.496, 88.58) = 24.57$, $p < .0001$] were observed (Figure S1A). During the period of peak effect (0-120 min post-dose), a significant linear trend was found [$F(1, 41) = 194.60$, $p < .0001$]; oxytocin dose-dependently reduced body temperature as dose increased (Figure S1B).

For the caffeine i.p. positive control experiment (Experiment 1b), no significant main effect of dose [$F(1, 12) = 0.06$, $p = .8069$] or interaction effect of dose x time [$F(1, 12) = 0.89$, $p = .3646$] were observed during baseline recording (pre-dose) for the last hour of the dark phase. Across the 6 hours following administration of i.p. caffeine, a significant main effect of dose [$F(1, 12) = 59.33$, $p < .0001$] and a significant interaction effect of dose x time [$F(5.799, 69.59) = 5.16$, $p = .0002$] were observed (Figure S1C). During the period of peak effect (0-120 min post-dose), caffeine significantly increased body temperature compared to VEH [$t(11) = 8.341$, $p < .0001$] (Figure S1D).

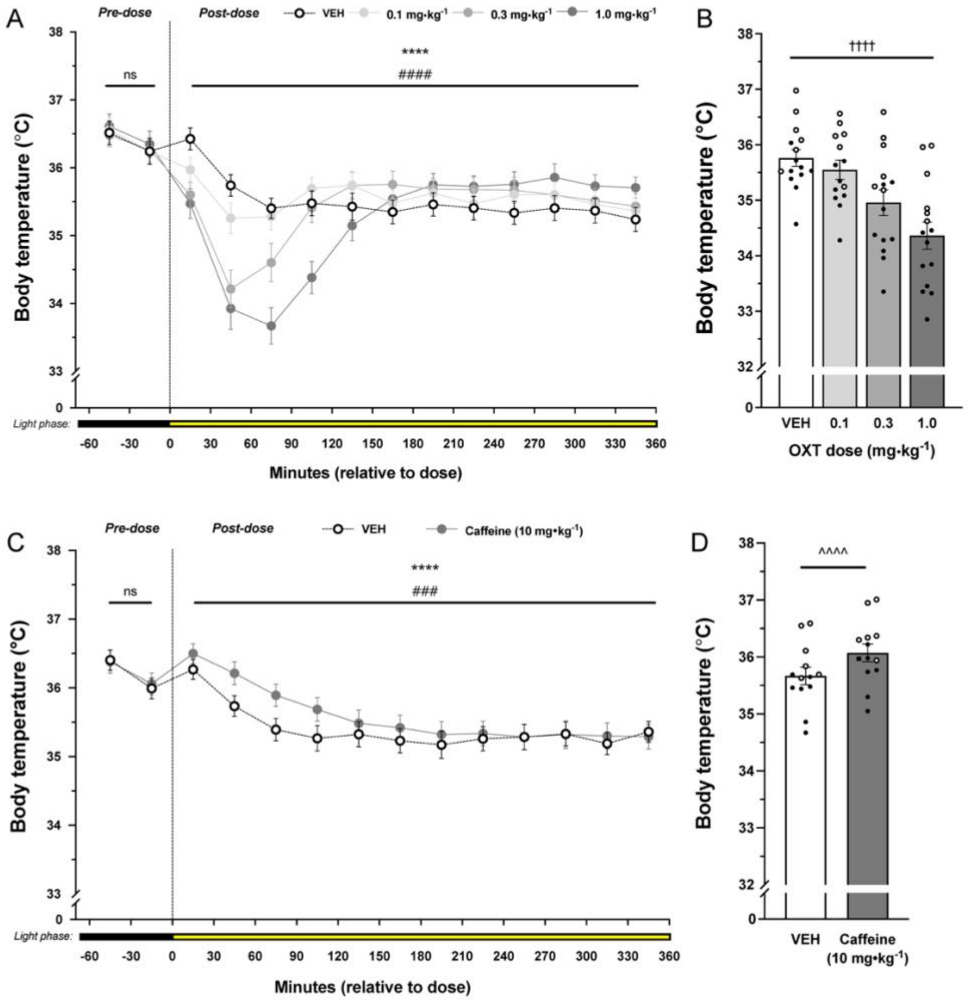

**Figure S1. Effects of i.p. oxytocin dose range and i.p. caffeine on body temperature.** Impact of oxytocin dose range (0, 0.1, 0.3, 1.0 mg·kg$^{-1}$) on body temperature (A) across the entire 7 hours of recording session (ZT23-6) and (B) aggregate value across period of peak effect (0-120 min, ZT0-2). Impact of caffeine (10 mg·kg$^{-1}$) on body temperature (C) across the entire 7 hours of recording session (ZT23-6) and (D) aggregate value across period of peak effect (0-120 min, ZT0-2). Data represent mean body temperature ± S.E.M., individual data points represent individual subject data, and the colour of each data point represents biological sex: white – male; black – female. Statistical test is indicated by the following symbols: * – dose main effect; # – dose x time interaction effect; † – linear trend; ^ – paired-sample t-test. Level of statistical significance is indicated by the number of symbols: one – $p < .05$, two – $p < .01$, three – $p < .001$, four – $p < .0001$.

*Baseline (1-hour dark phase pre-dose) data*

For the oxytocin i.p. dose response experiment (Experiment 1a), no significant effects were observed during the baseline period (last hour of dark phase) in any sleep-wake outcome assessed (Table S5).

**Table S5 – Sleep-wake outcomes at baseline in Experiment 1a.**

| Outcome | Dose (main effect) | p-value | Dose x Time interaction | p-value |
|---|---|---|---|---|
| ***Active wake*** | | | | |
| % | $F(2.701, 35.12) = 0.12$ | $p = .9432$ | $F(1.836, 22.64) = 0.36$ | $p = .6857$ |
| Bouts | $F(2.815, 36.59) = 1.94$ | $p = .1431$ | $F(2.064, 25.46) = 0.85$ | $p = .4417$ |
| Mean bout duration | $F(2.304, 29.95) = 1.13$ | $p = .3443$ | $F(2.375, 29.29) = 0.11$ | $p = .9246$ |
| ***Quiet wake*** | | | | |
| % | $F(2.566, 33.35) = 0.73$ | $p = .5233$ | $F(2.116, 26.09) = 0.04$ | $p = .9687$ |
| Bouts | $F(2.676, 34.79) = 1.78$ | $p = .1741$ | $F(2.049, 25.27) = 0.29$ | $p = .7575$ |
| Mean bout duration | $F(2.003, 26.04) = 0.48$ | $p = .6247$ | $F(1.502, 18.52) = 1.19$ | $p = .312$ |
| ***NREM sleep*** | | | | |
| % | $F(2.148, 27.93) = 0.55$ | $p = .5950$ | $F(1.707, 21.05) = 0.18$ | $p = .8049$ |
| Bouts | $F(2.391, 31.09) = 0.61$ | $p = .5796$ | $F(1.429, 17.62) = 0.38$ | $p = .6186$ |
| Mean bout duration | $F(2.243, 29.17) = 0.87$ | $p = .4405$ | $F(1.907, 23.52) = 0.70$ | $p = .4998$ |
| ***REM sleep*** | | | | |
| % | F(1.490, 19.37) = 0.42 | $p = .6018$ | $F(1.556, 19.20) = 0.63$ | $p = .5065$ |
| Bouts | F(1.873, 24.35) = 0.41 | $p = .6554$ | $F(1.897, 23.40) = 0.26$ | $p = .7602$ |
| Mean bout duration | F(2.477, 32.20) = 0.57 | $p = .6051$ | $F(2.065, 25.47) = 1.50$ | $p = .2413$ |

For the caffeine i.p. positive control experiment (Experiment 1b), during the baseline period (last hour of dark phase), VEH rats demonstrated longer mean duration of QW bouts than caffeine rats. Additionally, rats administered VEH demonstrated a significantly greater increase in %QW during the second-half of the baseline (pre-dose) period than the first-half compared to rats administered caffeine. No other significant effects were observed during the baseline period (last hour of dark phase) in sleep-wake outcome assessed in any sleep-wake outcome assessed (Table S6).

**Table S6 – Sleep-wake outcomes at baseline in Experiment 1b.**

| Outcome | Dose (main effect) | p-value | Dose x Time interaction | p-value |
|---|---|---|---|---|
| ***Active wake*** | | | | |
| % | $F$ (1, 11) = 0.07 | $p$ = .7960 | $F$ (1, 11) = 0.8010 | $p$ = .3899 |
| Bouts | $F$ (1, 11) = 0.74 | $p$ = .4077 | $F$ (1, 11) = 1.846 | $p$ = .2015 |
| Mean bout duration | $F$ (1, 11) = 0.18 | $p$ = .6771 | $F$ (1, 11) = 1.252 | $p$ = .2870 |
| ***Quiet wake*** | | | | |
| % | $F$ (1, 11) = 3.18 | $p$ = .1022 | $F$ (1, 11) = 5.34 | $p$ = .0412 |
| Bouts | $F$ (1, 11) = 0.36 | $p$ = .5584 | $F$ (1, 11) = 4.15 | $p$ = .0665 |
| Mean bout duration | $F$ (1, 11) = 11.55 | $p$ = .0059 | $F$ (1, 11) = 0.60 | $p$ = .4551 |
| ***NREM sleep*** | | | | |
| % | $F$ (1, 11) = 0.22 | $p$ = .6477 | $F$ (1, 11) < 0.01 | $p$ = .9711 |
| Bouts | $F$ (1, 11) = 0.36 | $p$ = .5591 | $F$ (1, 11) = 1.45 | $p$ = .2536 |
| Mean bout duration | $F$ (1, 11) = 3.45 | $p$ = .0901 | $F$ (1, 11) = 0.52 | $p$ = .4857 |
| ***REM sleep*** | | | | |
| % | $F$ (1, 11) = 0.15 | $p$ = .7063 | $F$ (1, 11) = 0.15 | $p$ = .7063 |
| Bouts | $F$ (1, 11) = 0.61 | $p$ = .4514 | $F$ (1, 11) = 0.22 | $p$ = .6486 |
| Mean bout duration | $F$ (1, 44) = 0.07 | $p$ = .7911 | $F$ (1, 44) = 0.14 | $p$ = .7145 |

#### *Experiment 2*

*Power spectral density data*

*Quiet wake*: No significant effects of any dose condition on ECoG PSD were detected at any frequency band during the first hour post-administration or the last 5 hours of recording.

*NREM sleep*: Relative to vehicle, oxytocin reduced ECoG PSD within the theta, sigma, and beta frequency bands, and these changes were not observed when L368,899 was administered prior to oxytocin.

*REM sleep*: Relative to vehicle, while oxytocin significantly increased ECoG PSD within the sigma frequency band, no effect of co-administration of oxytocin-L368,899 or L368,899 alone on PSD was found for any frequency band.

**Table S7 – Influence of oxytocin receptor antagonism on i.p. oxytocin (OXT) induced effects on power spectral density outcomes.**

| Relative to VEH | Antagonist-oxytocin dose condition | | |
|---|---|---|---|
| | ***VEH-OXT*** | ***L368,899-OXT*** | ***L368,899-VEH*** |
| ***QW (0-1 h)*** | | | |
| *Delta (0.1-4 Hz)* | $t(3) = 0.12$<br>$p = .9131$ | $t(3) = 0.43$<br>$p = .6983$ | $t(3) = 2.02$<br>$p = .1368$ |
| *Theta (4-8 Hz)* | $t(3) = 2.62$<br>$p = .0787$ | $t(3) = 0.04$<br>$p = .9705$ | $t(3) = 1.07$<br>$p = .3631$ |
| *Alpha (8-12 Hz)* | $t(3) = 1.02$<br>$p = .3847$ | $t(3) = 0.88$<br>$p = .4455$ | $t(3) = 0.35$<br>$p = .7467$ |
| *Sigma (12-16 Hz)* | $t(3) = 2.27$<br>$p = .1075$ | $t(3) = 0.09$<br>$p = .9330$ | $t(3) = 0.72$<br>$p = .5227$ |
| *Beta (16-25 Hz)* | $t(3) = 1.37$<br>$p = .2636$ | $t(3) = 0.38$<br>$p = .7286$ | $t(3) = 0.84$<br>$p = .4606$ |

| | | | |
|---|---|---|---|
| ***QW (1-6 h)*** | | | |
| *Delta (0.1-4 Hz)* | $t(3) = 0.37$<br>$p = .7385$ | $t(3) = 1.14$<br>$p = .3357$ | $t(3) = 0.10$<br>$p = .9270$ |
| *Theta (4-8 Hz)* | $t(3) = 0.99$<br>$p = .3960$ | $t(3) = 1.47$<br>$p = .2392$ | $t(3) = 0.20$<br>$p = .8510$ |
| *Alpha (8-12 Hz)* | $t(3) = 0.15$<br>$p = .8883$ | $t(3) = 0.78$<br>$p = .4948$ | $t(3) = 0.57$<br>$p = .6074$ |
| *Sigma (12-16 Hz)* | $t(3) = 0.39$<br>$p = .3542$ | $t(3) = 1.55$<br>$p = .5147$ | $t(3) = 2.64$<br>$p = .4299$ |
| *Beta (16-25 Hz)* | $t(3) = 0.83$<br>$p = .4686$ | $t(3) = 0.83$<br>$p = .4693$ | $t(3) = 0.46$<br>$p = .6791$ |
| ***NREM (0-1 h)*** | | | |
| *Delta (0.1-4 Hz)* | $t(3) = 1.80$<br>$p = .1704$ | $t(3) = 0.82$<br>$p = .4730$ | $t(3) = 1.10$<br>$p = .3502$ |
| *Theta (4-8 Hz)* | $t(3) = 3.31$<br>$p = .0453$ | $t(3) = 0.32$<br>$p = .7666$ | $t(3) = 2.28$<br>$p = .1069$ |
| *Alpha (8-12 Hz)* | $t(3) = 2.39$<br>$p = .0966$ | $t(3) = 0.44$<br>$p = .6908$ | $t(3) = 1.95$<br>$p = .1469$ |
| *Sigma (12-16 Hz)* | $t(3) = 3.50$<br>$p = .0396$ | $t(3) < 0.01$<br>$p = .9979$ | $t(3) = 2.96$<br>$p = .0594$ |
| *Beta (16-25 Hz)* | $t(3) = 3.26$<br>$p = .0472$ | $t(3) = 0.50$<br>$p = .6523$ | $t(3) = 1.68$<br>$p = .1922$ |
| ***NREM (1-6 h)*** | | | |
| *Delta (0.1-4 Hz)* | $t(3) = 0.54$<br>$p = .6258$ | $t(3) = 1.18$<br>$p = .3233$ | $t(3) = 0.23$<br>$p = .8300$ |
| *Theta (4-8 Hz)* | $t(3) = 1.10$<br>$p = .3520$ | $t(3) = 0.91$<br>$p = .4306$ | $t(3) = 0.59$<br>$p = .5975$ |
| *Alpha (8-12 Hz)* | $t(3) = 0.17$<br>$p = .8787$ | $t(3) = 0.73$<br>$p = .5176$ | $t(3) = 0.03$<br>$p = .9746$ |
| *Sigma (12-16 Hz)* | $t(3) = 0.08$<br>$p = .9429$ | $t(3) = 0.12$<br>$p = .9145$ | $t(3) = 0.60$<br>$p = .5968$ |
| *Beta (16-25 Hz)* | $t(3) < 0.01$<br>$p = .9985$ | $t(3) = 0.77$<br>$p = .4951$ | $t(3) = 0.05$<br>$p = .9652$ |
| ***REM (0-6 h)*** | | | |
| *Delta (0.1-4 Hz)* | $t(3) = 0.48$<br>$p = .6655$ | $t(3) = 0.97$<br>$p = .4019$ | $t(3) = 0.35$<br>$p = .7508$ |
| *Theta (4-8 Hz)* | $t(3) = 0.02$<br>$p = .9880$ | $t(3) = 0.76$<br>$p = .5020$ | $t(3) = 1.01$<br>$p = .3883$ |
| *Alpha (8-12 Hz)* | $t(3) = 0.65$<br>$p = .5606$ | $t(3) = 0.93$<br>$p = .4224$ | $t(3) = 0.35$<br>$p = .7510$ |
| *Sigma (12-16 Hz)* | $t(3) = 11.04$<br>$p = .0016$ | $t(3) = 1.46$<br>$p = .2399$ | $t(3) = 1.65$<br>$p = .1973$ |
| *Beta (16-25 Hz)* | $t(3) = 2.40$<br>$p = .0959$ | $t(3) = 1.07$<br>$p = .3623$ | $t(3) = 1.25$<br>$p = .3009$ |

### *Oestrus data*

No significant differences in the proportion of rats in proestrus/oestrus and metoestrus/dioestrus stages between dose conditions and their respective VEH controls were observed in Experiment 2 (Table S8).

**Table S8 – Proportion of rats in oestrus phases for each dose condition in Experiment 2.**

| Dose condition | Proestrus/Oestrus (%) | Metoestrus/Dioestrus (%) | Fisher's exact test (2-sided) |
|---|---|---|---|
| **VEH-VEH** | 25 | 75 | |
| **VEH-OXT** | 50 | 50 | $p > .9999$[a] |
| **L368,899-VEH** | 75 | 25 | $p > .9999$[b] |
| **L368,899-OXT** | 50 | 50 | $p > .9999$[c] |

[a]VEH-VEH vs VEH-OXT
[b]L368,899-VEH vs L368,899-OXT
[c]VEH-OXT vs L368,899-OXT

*Body temperature data*

No significant dose x antagonist interaction effect was observed [$F(1,3) = 6.91$, $p = .0785$] during the period of peak effect from 0-120 min post-dose (Figure S2). However, pairwise comparisons revealed that 1 mg·kg$^{-1}$ oxytocin significantly reduced body temperature—VEH-oxytocin vs VEH-VEH [$t(3) = 18.08$, $p = .0004$]— and that pre-administration of L368,899 significantly attenuated the oxytocin-induced reduction on body temperature—VEH-oxytocin vs L368,899-oxytocin [$t(3) = 3.465$, $p = .0405$].

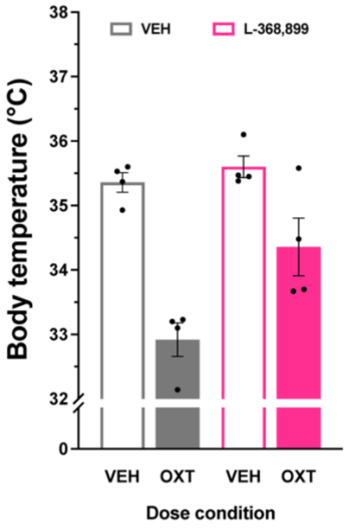


**Figure S2. Effects of oxytocin receptor antagonism of i.p. oxytocin-induced reduction in body temperature.** Impact of oxytocin receptor antagonism using L368,899 (i.p., 5 mg·kg$^{-1}$) on the 1 mg·kg$^{-1}$ (i.p.) oxytocin-induced reduction in body temperature in female rats. Data represent mean body temperature ± S.E.M. for aggregate value across the period of peak effect (0-120 min post-dose, ZT0-2), and individual data points represent individual subject data. A significant *Dose* x *Antagonist* interaction effect is represented by * and level of statistical significance is indicated by the number of symbols: * – $p < .05$.

***Experiment 3***

*Power spectral density data*

*Quiet wake*: No effects of i.n. oxytocin on ECoG PSD were observed at any dose at any time. During the last 5 hours of recording, i.n. caffeine significantly reduced ECoG PSD relative to vehicle in the sigma and beta frequency bands, however no other significant effects of caffeine on ECoG PSD outcomes were found.

*NREM sleep*: No effects of i.n. oxytocin on ECoG PSD were observed at any dose at any time. During the first hour post-administration, caffeine significantly reduced ECoG PSD within the delta, theta, alpha and sigma frequency bands, however no effect of caffeine was detected for the following 5 hours of recording.

*REM sleep*: No effects of i.n. oxytocin or i.n. caffeine on ECoG PSD were observed at any dose at any time.

**Table S9 – Effects of i.n. oxytocin dose range and i.n. caffeine dose on power spectral density outcomes during quiet wake, NREM sleep and REM sleep.**

| **Relative to VEH** | **Oxytocin (mg·kg$^{-1}$)** | | | **Caffeine (mg·kg$^{-1}$)** |
|---|---|---|---|---|
| | ***0.06*** | ***1.0*** | ***3.0*** | ***10*** |
| ***QW (0-1 h)*** | | | | |
| *Delta (0.1-4 Hz)* | $t(5) = 0.05$ $p = .9651$ | $t(5) = 0.24$ $p = .8216$ | $t(3) = 0.03$ $p = .9770$ | $t(4) = 0.31$ $p = .7725$ |
| *Theta (4-8 Hz)* | $t(5) = 0.47$ $p = .6611$ | $t(5) = 0.58$ $p = .5864$ | $t(3) = 1.25$ $p = .2998$ | $t(4) = 0.39$ $p = .7306$ |
| *Alpha (8-12 Hz)* | $t(5) = 0.14$ $p = .8921$ | $t(5) = 0.16$ $p = .8815$ | $t(3) = 0.34$ $p = .7582$ | $t(4) = 0.06$ $p = .9534$ |
| *Sigma (12-16 Hz)* | $t(5) = 0.99$ $p = .3691$ | $t(5) = 0.37$ $p = .7234$ | $t(3) = 0.97$ $p = .4028$ | $t(4) = 1.79$ $p = .1483$ |
| *Beta (16-25 Hz)* | $t(5) = 0.03$ $p = .9768$ | $t(5) = 0.49$ $p = .6479$ | $t(3) = 0.68$ $p = .5438$ | $t(4) = 1.77$ $p = .1509$ |
| ***QW (1-6 h)*** | | | | |
| *Delta (0.1-4 Hz)* | $t(5) = 0.20$ $p = .8479$ | $t(5) = 0.24$ $p = .8187$ | $t(3) = 0.09$ $p = .9332$ | $t(4) = 0.36$ $p = .7391$ |
| *Theta (4-8 Hz)* | $t(5) = 0.15$ $p = .8870$ | $t(5) = 0.44$ $p = .6789$ | $t(3) = 0.37$ $p = .7250$ | $t(4) = 0.05$ $p = .9613$ |
| *Alpha (8-12 Hz)* | $t(5) = 0.24$ $p = .8205$ | $t(5) = 0.35$ $p = .7399$ | $t(3) = 0.53$ $p = .6312$ | $t(4) = 0.26$ $p = .8042$ |
| *Sigma (12-16 Hz)* | $t(5) = 0.56$ $p = .6001$ | $t(5) = 0.19$ $p = .8576$ | $t(3) = 1.20$ $p = .3160$ | $t(4) = 2.90$ $p = .0440$ |
| *Beta (16-25 Hz)* | $t(5) = 0.97$ $p = .3772$ | $t(5) = 0.18$ $p = .8609$ | $t(3) = 0.60$ $p = .5885$ | $t(4) = 4.77$ $p = .0088$ |
| ***NREM (0-1 h)*** | | | | |
| *Delta (0.1-4 Hz)* | $t(5) = 0.03$ $p = .9808$ | $t(5) = 0.34$ $p = .7488$ | $t(3) = 0.02$ $p = .9830$ | $t(3) = 3.41$ $p = .0423$ |
| *Theta (4-8 Hz)* | $t(5) = 0.05$ $p = .9595$ | $t(5) = 0.37$ $p = .7281$ | $t(3) = 0.30$ $p = .7810$ | $t(3) = 4.13$ $p = .0257$ |
| *Alpha (8-12 Hz)* | $t(5) = 0.26$ $p = .8084$ | $t(5) = 0.03$ $p = .9807$ | $t(3) = 0.32$ $p = .7690$ | $t(3) = 3.89$ $p = .0302$ |
| *Sigma (12-16 Hz)* | $t(5) = 0.43$ $p = .6830$ | $t(5) = 0.53$ $p = .6204$ | $t(3) = 0.15$ $p = .8876$ | $t(3) = 5.39$ $p = .0125$ |
| *Beta (16-25 Hz)* | $t(5) = 0.72$ $p = .5039$ | $t(5) = 0.06$ $p = .9562$ | $t(3) = 0.31$ $p = .7739$ | $t(3) = 3.05$ $p = .0552$ |
| ***NREM (1-6 h)*** | | | | |
| *Delta (0.1-4 Hz)* | $t(5) = 0.25$ $p = .8148$ | $t(5) = 0.29$ $p = .7809$ | $t(3) = 0.06$ $p = .9558$ | $t(4) = 0.38$ $p = .7212$ |
| *Theta (4-8 Hz)* | $t(5) = 0.37$ $p = .7272$ | $t(5) = 0.20$ $p = .8479$ | $t(3) = 0.33$ $p = .7585$ | $t(4) = 0.32$ $p = .7681$ |
| *Alpha (8-12 Hz)* | $t(5) = 0.35$ $p = .7416$ | $t(5) = 0.01$ $p = .9896$ | $t(3) = 0.19$ $p = .8612$ | $t(4) = 0.13$ $p = .9004$ |
| *Sigma (12-16 Hz)* | $t(5) = 1.16$ $p = .2978$ | $t(5) = 0.45$ $p = .6737$ | $t(3) = 0.01$ $p = .9890$ | $t(4) = 0.10$ $p = .9278$ |
| *Beta (16-25 Hz)* | $t(5) = 0.64$ $p = .5523$ | $t(5) = 0.04$ $p = .9666$ | $t(3) = 0.28$ $p = .7943$ | $t(4) = 0.07$ $p = .9492$ |
| ***REM (0-6 h)*** | | | | |
| *Delta (0.1-4 Hz)* | $t(5) < 0.01$ $p = .9974$ | $t(5) = 0.30$ $p = .7748$ | $t(3) = 0.08$ $p = .9422$ | $t(5) = 0.25$ $p = .8113$ |
| *Theta (4-8 Hz)* | $t(5) = 0.10$ $p = .9267$ | $t(5) = 0.12$ $p = .9074$ | $t(3) = 0.07$ $p = .9485$ | $t(5) = 0.20$ $p = .8502$ |
| *Alpha (8-12 Hz)* | $t(5) = 0.01$ $p = .9889$ | $t(5) = 0.08$ $p = .9374$ | $t(3) = 0.15$ $p = .8892$ | $t(5) = 0.36$ $p = .7321$ |
| *Sigma (12-16 Hz)* | $t(5) = 0.06$ $p = .9577$ | $t(5) = 0.32$ $p = .7594$ | $t(3) = 0.12$ $p = .9157$ | $t(5) = 0.18$ $p = .8624$ |
| *Beta (16-25 Hz)* | $t(5) = 0.66$ $p = .5360$ | $t(5) = 0.47$ $p = .6582$ | $t(3) = 1.11$ $p = .3496$ | $t(5) = 0.17$ $p = .8754$ |

#### *Oestrus data*

No significant differences were observed in the proportion of rats in proestrus/oestrus and metoestrus/dioestrus stages between oxytocin dose conditions in Experiment 3a (Table S10).

Additionally, while no significant difference in the proportion of rats in proestrus/oestrus and metoestrus/dioestrus stages between the 3 mg·kg$^{-1}$ dose and VEH conditions, a significant difference was observed in these proportions between 10 mg·kg$^{-1}$ caffeine and VEH conditions in Experiment 3b (Table S11).

**Table S10. Proportion of rats in oestrus phases for each dose condition in Experiment 3a.**

| OXT dose (mg·kg$^{-1}$) | Proestrus/Oestrus (%) | Metoestrus/Dioestrus (%) | Fisher's exact test (2-sided) |
|---|---|---|---|
| **VEH** | 66.7 | 33.3 | |
| **0.06** | 33.33 | 66.7 | $p$ = .5671 |
| **1** | 33.33 | 66.7 | $p$ = .5671 |

**Table S11. Proportion of rats in oestrus phases for each dose condition in Experiment 3b.**

| Dose condition | Proestrus/Oestrus (%) | Metoestrus/Dioestrus (%) | Fisher's exact test (2-sided) |
|---|---|---|---|
| **VEH** | 80 | 20 | |
| **OXT (3 mg·kg$^{-1}$)** | 50 | 50 | $p$ = .5238 |
| **Caffeine (10 mg·kg$^{-1}$)** | 0 | 100 | $p$ = .0476 |

*Body temperature data*

For the oxytocin i.n. dose response experiment (Experiment 3a), no significant main effect of dose [$F$(1.804, 9.021) = 0.83, $p$ = .4562] or interaction effect of dose x time [$F$(1.649, 8.245) = 0.44, $p$ = .6201] were observed during baseline recording (pre-dose) for the last hour of the dark phase. Across the 6 hours post-administration of i.n. oxytocin, no significant main effect of dose [$F$(1.616, 8.081) = 0.67, $p$ = .5089] or interaction effect of dose x time [$F$(3.550, 17.75) = 0.62, $p$ = .6371] were observed (Figure S3A). During the period of peak effect (0-120 min post-dose), no significant linear trend was found [$F$(1, 15) = 0.94, $p$ = .3481] (Figure S3B).

For Experiment 3b assessing the effects of i.n. oxytocin (3 mg·kg$^{-1}$) and caffeine (10 mg·kg$^{-1}$), averaged over time, no significant effect of oxytocin [$MD$ = 0.47, S.E.M. = 0.31, $t$(3) = 1.52, $p$ = .2260] or caffeine [$MD$ = 0.55, S.E.M. = 0.27, $t$(4) = 2.05, $p$ = .1104] were observed during baseline recording (pre-dose) for the last hour of the dark phase. Averaged over the 6 hours post i.n. dose, no significant main effect of oxytocin [$MD$ = 0.27, S.E.M. = 0.45, $t$(3) = .59, $p$ = .5942] and a significant interaction effect of dose x time [$MD$ = 0.35, 0.37, t(4) = .95, $p$ = .3974] were observed (Figure S3C). During the period of peak effect (0-120 min post-dose), no significant pairwise comparisons were found between oxytocin vs VEH [$MD$ = 0.18, S.E.M. = 0.57, $t$(3) = .32, $p$ = .7708] or between caffeine and VEH [$MD$ = 0.22, S.E.M. = 0.40, $t$(4) = .54, $p$ = .6177] (Figure S3D).

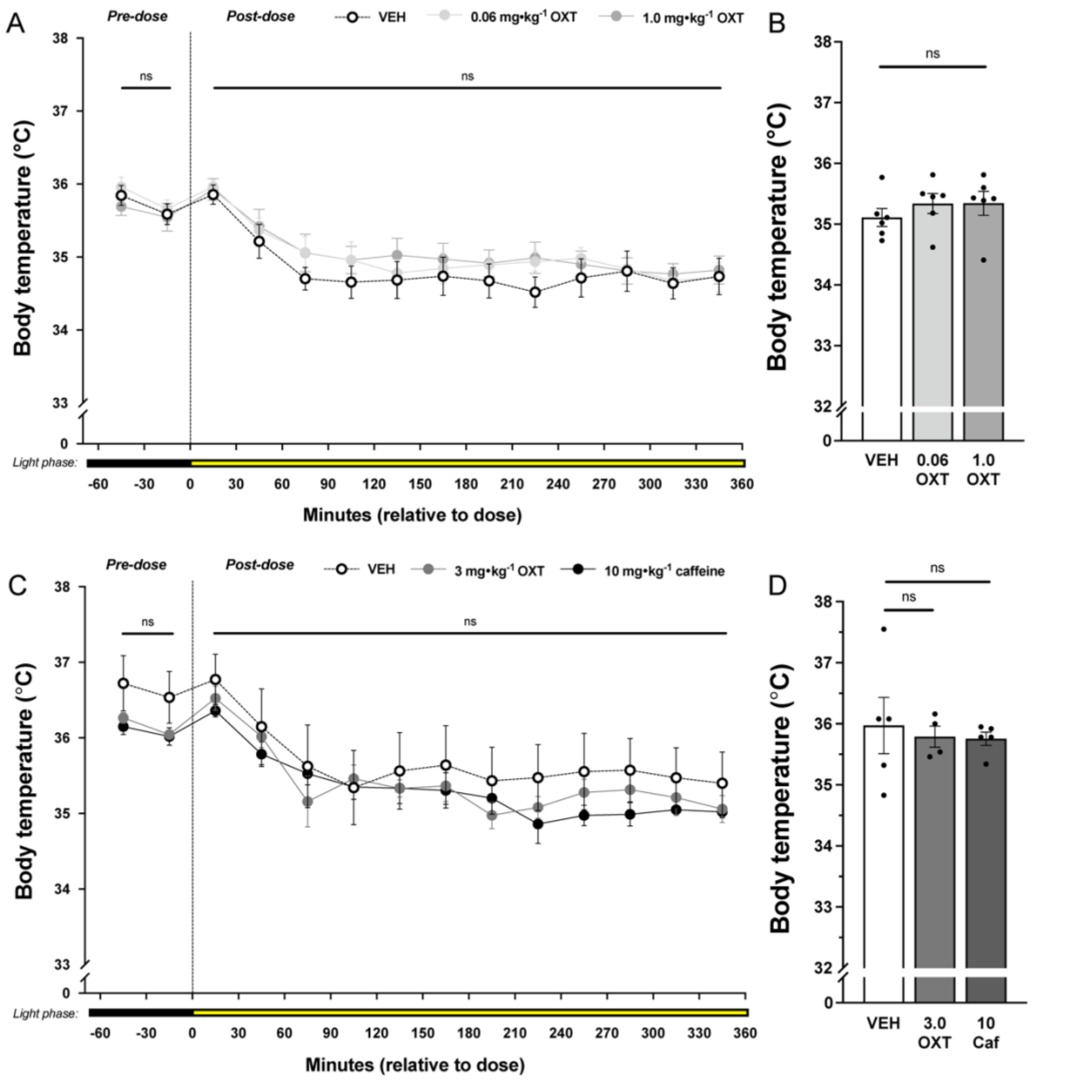


**Figure S3. Effects of i.n. oxytocin dose range and i.n. caffeine on body temperature.** Impact of i.n. oxytocin dose range (0, 0.06, 1.0 mg·kg$^{-1}$) on body temperature (A) across the entire 7 hours of recording session (ZT23-6) and (B) aggregate value across period of peak effect (0-120 min, ZT0-2). Impact of i.n. oxytocin (3 mg·kg$^{-1}$) caffeine (10 mg·kg$^{-1}$) on body temperature (C) across the entire 7 hours of recording session (ZT23-6) and (D) aggregate value across period of peak effect (0-120 min, ZT0-2). Data represent mean body temperature ± S.E.M. and individual data points represent individual subject data. Statistical test is indicated by the following symbols: * – dose main effect; # – dose x time interaction effect; † – linear trend; ^ – paired-sample t-test. Level of statistical significance is indicated by the number of symbols: * $p < .05$, ** $p < .01$, *** $p < .001$, **** $p < .0001$.

### *Baseline (1-hour dark phase) data*

For the oxytocin i.n. dose response experiment (Experiment 3a), no significant effects were observed during the baseline period (last hour of dark phase) on any sleep-wake outcome assessed (Table S12).

**Table S12 – Sleep-wake outcomes at baseline in Experiment 3a.**

| **Outcome** | **Dose (main effect)** | **p-value** | **Dose x Time interaction** | **p-value** |
|---|---|---|---|---|
| ***Active wake*** | | | | |

| | | | | |
|---|---|---|---|---|
| % | *F*(1.088, 5.439) = 2.10 | *p* = .2046 | *F*(1.278, 6.390) = 0.57 | *p* = .5201 |
| Bouts | *F*(1.802, 9.008) = 0.18 | *p* = .8160 | *F*(1.995, 9.975) = 0.26 | *p* = .7772 |
| Mean bout duration | *F*(1.140, 5.700) = 2.41 | *p* = .1754 | *F*(1.881, 9.404) = 1.37 | *p* = .2987 |
| ***Quiet wake*** | | | | |
| % | *F*(1.874, 9.368) = 0.11 | *p* = .8854 | *F*(1.568, 7.840) = 0.38 | *p* = .646 |
| Bouts | *F*(1.954, 9.768) = 0.77 | *p* = .4869 | *F*(1.388, 6.939) = 0.28 | *p* = .6866 |
| Mean bout duration | *F*(1.347, 6.733) = 0.05 | *p* = .8992 | *F*(1.112, 5.560) = 0.14 | *p* = .7500 |
| ***NREM sleep*** | | | | |
| % | *F*(1.141, 5.706) = 1.07 | *p* = .3553 | *F*(1.340, 6.698) = 0.38 | *p* = .6193 |
| Bouts | *F*(1.626, 8.130) = 1.46 | *p* = .2816 | *F*(1.909, 9.543) = 2.77 | *p* = .1141 |
| Mean bout duration | *F*(1.739, 8.693) = 0.45 | *p* = .6270 | *F*(1.440, 7.200) = 0.02 | *p* = .9579 |
| ***REM sleep*** | | | | |
| % | *F*(1.925, 9.627) = 2.97 | *p* = .1003 | *F*(1.842, 9.212) = 3.18 | *p* = .0918 |
| Bouts | *F*(1.668, 8.338) = 2.99 | *p* = .1099 | *F*(1.871, 9.357) = 3.61 | *p* = .0710 |
| Mean bout duration | *F*(1.483, 22.24) = 0.56 | *p* = .5295 | *F*(1.416, 21.24) = 0.66 | *p* = .4759 |

For the Experiment 3b, assessing the impact of 3 mg·kg$^{-1}$ i.n. oxytocin and 10 mg·kg$^{-1}$ caffeine i.n., no significant effects were observed during the baseline period (last hour of dark phase) on any sleep-wake outcome assessed (Table S13).

**Table S13 – Sleep-wake outcomes at baseline in Experiment 3b.**

| **Outcome** | **VEH vs OXT 3 mg·kg$^{-1}$** ***MD*, SEM, *t*** | **p-value** | **VEH vs Caffeine 10 mg·kg$^{-1}$** ***MD*, SEM, *t*** | **p-value** |
|---|---|---|---|---|
| ***Active wake*** | | | | |
| % | -10.07, 7.993, *t*(3) = 1.26 | *p* = .2967 | -9.6, 15.42, *t*(4) = 0.62 | *p* = .5673 |
| Bouts | 0.6, 3.212, *t*(3) = 0.19 | *p* = .8637 | -1.9, 1.764, *t*(4) = 1.08 | *p* = .3419 |
| Mean bout duration | -25.13, 20.89, *t*(3) = 1.20 | *p* = .3153 | -7.22, 12.04, *t*(4) = 0.60 | *p* = .5809 |
| ***Quiet wake*** | | | | |
| % | 2.988, 6.226, *t*(3) = 0.48 | *p* = .6642 | 3.180, 2.170, *t*(4) = 1.47 | *p* = .2167 |
| Bouts | 3.725, 3.335, *t*(3) = 1.12 | *p* = .3454 | 1, 0.866, *t*(4) = 1.16 | *p* = .3125 |
| Mean bout duration | 0.225, 3.591, *t*(3) = 0.06 | *p* = .9540 | 1.29, 1.665, *t*(4) = 0.77 | *p* = .4818 |
| ***NREM sleep*** | | | | |
| % | 6.070, 3.014, *t*(3) = 2.01 | *p* = .1375 | 6.290, 13.90, *t*(4) = 0.45 | *p* = .6743 |
| Bouts | 1.725, 1.818, *t*(3) = 0.95 | *p* = .4126 | 1.6, 2.015, *t*(4) = 0.79 | *p* = .4716 |
| Mean bout duration | 15.08, 14.69, *t*(3) = 1.03 | *p* = .3800 | 4.93, 21.89, *t*(4) = 0.23 | *p* = .8329 |
| ***REM sleep*** | | | | |
| % | 1, 0.8179, *t*(3) = 1.22 | *p* = .3088 | 0.89, 0.6972, *t*(4) = 1.28 | *p* = .2708 |
| Bouts | 0.5, 0.3623, *t*(3) = 1.38 | *p* = .2614 | 0.3, 0.3, *t*(4) = 1.00 | *p* = .3739 |
| Mean bout duration | 6.83, 5.087, *t*(3) = 1.34 | *p* = .2720 | 5.83, 4.213, *t*(4) = 1.38 | *p* = .2386 |
